# Eliminating accidental deviations to minimize generalization error and maximize replicability: applications in connectomics and genomics

**DOI:** 10.1101/802629

**Authors:** Eric W. Bridgeford, Shangsi Wang, Zhi Yang, Zeyi Wang, Ting Xu, Cameron Craddock, Jayanta Dey, Gregory Kiar, William Gray-Roncal, Carlo Colantuoni, Christopher Douville, Stephanie Noble, Carey E. Priebe, Brian Caffo, Michael Milham, Xi-Nian Zuo, Consortium for Reliability and Reproducibility, Joshua T. Vogelstein

## Abstract

Replicability, the ability to replicate scientific findings, is a prerequisite for scientific discovery and clinical utility. Troublingly, we are in the midst of a replicability crisis. A key to replicability is that multiple measurements of the same item (e.g., experimental sample or clinical participant) under fixed experimental constraints are relatively similar to one another. Thus, statistics that quantify the relative contributions of accidental deviations—such as measurement error—as compared to systematic deviations—such as individual differences—are critical. We demonstrate that existing replicability statistics, such as intra-class correlation coefficient and fingerprinting, fail to adequately differentiate between accidental and systematic deviations in very simple settings. We therefore propose a novel statistic, *discriminability*, which quantifies the degree to which an individual’s samples are relatively similar to one another, without restricting the data to be univariate, Gaussian, or even Euclidean. Using this statistic, we introduce the possibility of optimizing experimental design via increasing discriminability and prove that optimizing discriminability improves performance bounds in subsequent inference tasks. In extensive simulated and real datasets (focusing on brain imaging and demonstrating on genomics), only optimizing data *discriminability* improves performance on all subsequent inference tasks for each dataset. We therefore suggest that designing experiments and analyses to optimize discriminability may be a crucial step in solving the replicability crisis, and more generally, mitigating accidental measurement error.

**Author Summary:** In recent decades, the size and complexity of data has grown exponentially. Unfortunately, the increased scale of modern datasets brings many new challenges. At present, we are in the midst of a replicability crisis, in which scientific discoveries fail to *replicate* to new datasets. Difficulties in the measurement procedure and measurement processing pipelines coupled with the influx of complex high-resolution measurements, we believe, are at the core of the replicability crisis. If measurements themselves are not replicable, what hope can we have that we will be able to use the measurements for replicable scientific findings? We introduce the “discriminability” statistic, which quantifies how *discriminable* measurements are from one another, without limitations on the structure of the underlying measurements. We prove that discriminable strategies tend to be strategies which provide better accuracy on downstream scientific questions. We demonstrate the utility of discriminability over competing approaches in this context on two disparate datasets from both neuroimaging and genomics. Together, we believe these results suggest the value of designing experimental protocols and analysis procedures which optimize the discriminability.

## 1 Introduction

Understanding variability, and the sources thereof, is fundamental to all of data science. Even the first papers on modern statistical methods concerned themselves with distinguishing accidental from systematic variability [1]. Accidental deviations correspond to sources of variance that are not of scientific interest, including measurement noise and artefacts from the particular experiment (often called “batch effects” [2]). Quantifying systematic deviations of the variables of interest, such as variance across items within a study, is often the actual goal of the study. Thus, delineating between these two sources of noise is a central quest in data science, and failure to do so, has been problematic in modern science [3].

Scientific replicability, or the degree to which a result can be replicated using the same methods applied to the same scientific question on new data [4], is key in data science, whether applied to basic discovery or clinical utility [5]. As a rule, if results do not replicate, we can not justifiably trust them [4] (though replication does not imply validation necessarily [6]). The concept of replicability is closely related to the statistical concepts of stability [7] and robustness [5]. Engineering and operations research have been concerned with *reliability* for a long time, as they require that their products are reliable under various conditions. Very recently, the general research community became interested in these issues, as individuals began noticing and publishing failures to replicate across fields, including neuroscience and psychology [8–10].

A number of strategies have been suggested to resolve this “replicability crisis.” For example, the editors of “Basic and Applied Social Psychology” have banned the use of p-values [11]. Unfortunately, an analysis of the publications since banning indicates that studies after the ban tended to overstate, rather than understate, their claims, suggesting that this proposal possibly had the opposite effect [12]. More recently, the American Statistical Association released a statement recommending banning the phrase “statistically significant” for similar reasons [13, 14].

A different strategy has been to quantify the repeatability of one’s measurements by measuring each sample (or individual) multiple times. Such “test-retest reliability” experiments quantify the relative similarity of multiple measurements of the same item, as compared to different items [15]. Approaches which investigate *measurement repeatability* quantify the degree to which measurements obtained in one session are similar to a set of measurements obtained in a second session, to test replicability [4]. This practice has been particularly popular in brain imaging, where many studies have been devoted to quantifying the repeatability of different univariate properties of the data [16–19]. In practice, however, these approaches have severe limitations. The Intraclass Correlation Coefficient (`ICC`) is an approach that quantifies the ratio of within item variance to across item variance. The `ICC` is univariate, with limited applicability to high-dimensional data, and its interpretation suffers from limitations due to its motivating Gaussian assumptions. Previously proposed generalizations of `ICC`, such as the Image Intraclass Correlation Coefficient (`I2C2`), generalize `ICC` to multivariate data, but require large sample sizes to estimate high-dimensional covariance matrices. Further, motivating intuition of `I2C2` makes similar Gaussian parametric assumptions as `ICC`, and therefore exhibits similar limitations. The Fingerprinting Index (`Fingerprint`) provides a nonparametric and multivariate technique for quantifying test-retest reliability, but its greedy assignment leads it to provide counter-intuitive results in certain contexts. A number of other approaches such as `NPAIRS` [20, 21] provide general frameworks for evaluating activation-based neuroimaging timeseries experiments, which can be extended to other modalities [22, 23]. A thorough discussion and analysis of these and similar approaches is provided in Supporting Information S1.

Perhaps the most problematic aspect of these approaches is clear from the popular adage, “garbage in, garbage out” [24]. If the measurements themselves are not sufficiently replicable, then scalar summaries of the data cannot be replicable either. This primacy of measurement is fundamental in statistics, so much so that one of the first modern statistics textbook, R.A. Fisher’s, “The Design of Experiments” [25], is focused on taking measurements. Motivated by Fisher’s work on experimental design, and Spearman’s work on measurement, rather than recommending different post-data acquisition inferential techniques, or computing the repeatability of data after collecting, we take a different approach. Specifically, **we advocate for explicitly and specifically designing experiments to ensure that they provide highly replicable data, rather than hoping that they do and performing post-hoc checks after collecting the data**. Thus, we concretely recommend that new studies leverage existing protocols that have previously been established to generate highly replicable data. If no such protocols are available for your question, we recommend designing new protocols in such a way that replicability is explicitly considered (and not compromised) in each step of the design. Experimental design has a rich history, including in psychology [26] and neuroscience [27, 28]. The vast majority of work in experimental design, however, focuses on designing an experiment to answer a particular scientific question. In this big data age, experiments are often designed to answer many questions, including questions not even considered at the time of data acquisition. How can one even conceivably design experiments to obtain data that is particularly useful for those questions?

We propose to design experiments to optimize the *inter-item discriminability* of individual items (for example, participants in a study, or samples in an experiment). This idea is closely inspired by and related to ideas proposed by Cronbach’s “Theory of Generalizability” [29, 30]. To do so, we leverage our recently introduced `Discr` statistic [31]. `Discr` quantifies the degree to which multiple measurements of the same item are more similar to one another than they are to other items [31], essentially capturing the desiderata of Spearman from over 100 years ago. This statistic has several advantages over existing statistics that one could potentially use to optimize experimental design. First, it is nonparametric, meaning that its validity and interpretation do not depend on any parametric assumptions, such as Gaussianity. Second, it can readily be applied to multivariate Euclidean data, or even non-Euclidean data (such as images, text, speech, or networks). Third, it can be applied to any stage of the data science pipeline, from data acquisition to data wrangling to data inferences. Finally, and most uniquely, one of the main advantages of `ICC`, is that under certain assumptions, `ICC` can provide an upper bound on predictive accuracy for any subsequent inference task. Specifically, we present here a result generalizing `ICC`’s bound on predictive accuracy to a multivariate additive noise setting. Thus, `Discr` is the *only* non-parametric multivariate measure of test-retest reliability with formal theoretical guarantees of convergence and upper bounds on subsequent inference performance. We show that this property makes `Discr` desirable through empirical simulations and across multiple scientific domains. An important clarification is that high test-retest reliability does not provide any information about the extent to which a measurement coincides with what it is purportedly measuring (construct validity). Even though replicable data are not enough on their own, replicable data are required for stable subsequent inferences.

This manuscript provides the following contributions:

1. Demonstrates that `Discr` is a statistic that adequately quantifies the relative contribution of certain accidental and systematic deviations, whereas previously proposed statistics have not.
2. Formalizes hypothesis tests to assess discriminability of a dataset, and whether one dataset or approach is more discriminable than another. This is in contrast to previously proposed non-parametric approaches to quantify test-retest reliability, that merely provide a test statistic, but no valid test per se.
3. Provides sufficient conditions for `Discr` to provide a lower bound on predictive accuracy. `Discr` is the *only* multivariate measure of replicability that has been theoretically related to criterion validity.
4. Illustrates on 28 neuroimaging datasets from Consortium for Reliability and Reproducibility (CoRR) [32] and two genomics datasets (i) the preprocessing pipelines which maximize `Discr`, and (ii) that maximizing `Discr` is significantly associated with maximizing the amount of information about multiple covariates, in contrast to other related statistics.
5. Provides all source code and data derivatives open access at https://neurodata.io/mgc.

## 2 Methods

### 2.1 The inter-item discriminability statistic

Testing for inter-item discriminability is closely related to, but distinct from, k-sample testing. In k-sample testing we observe k groups, and we want to determine whether they are different *at all*. In inter-item discriminability, the k groups are in fact k different items (or individuals), and we care about whether replicates within each of the k groups are close to each other, which is a specific kind of difference. As a general rule, if one can specify the kind of difference one is looking for, then tests can have more power for that particular kind of difference. The canonical example of this would be an t-test, where if only looks at whether the means are different across the groups, one obtains higher power than if also looking for differences in variances.

To give a concrete example, assume one item has replicates on a circle with radius one, with random angles. Consider another item whose replicates live on another circle, concentric with the first, but with a different radius. The two items differ, and many nonparametric two-sample tests would indicate so (because one can perfectly identify the item by the radius of the sample). However, the discriminability in this example is not one, because there are samples of either item that are further from other samples of that item than samples from the other item.

On this basis, we developed our inter-item discriminability test statistic (`Discr`), which is inspired by, and builds upon, nonparametric two-sample and k-sample testing approaches called “Energy statistics” [33] and “Kernel mean embeddings” [34] (which are equivalent [35]). These approaches compute all pairwise similarities (or distances) and operate on them. `Discr` differs from these methods in two key ways. First, rather than operating on the magnitudes of all the pairwise distances directly, `Discr` operates on the ranks of the distances, rendering it robust to monotonic transformations of the data [36]. Second, `Discr` only considers comparisons of the ranks of pairwise distances between different items with the ranks of pairwise distances between the same item. All other information is literally discarded, as it does not provide insight into the question of interest.

Fig 1 shows three different simulations illustrating the differences between `Discr` and other replicability statistics, including the fingerprinting index (`Fingerprint`) [37], intraclass correlation coefficient (`ICC`) [38], and `Kernel` [34] (see Supporting Information S1 for details). All four statistics operate on the pairwise distance matrices in column (B). However, `Discr`, unlike the other statistics, only considers the elements of each row whose magnitudes are smaller than the distances within an item. Thus, `Discr` explicitly quantifies the degree to which multiple measurements of the same item are more similar to one another than they are to other items.

**Fig 1.**
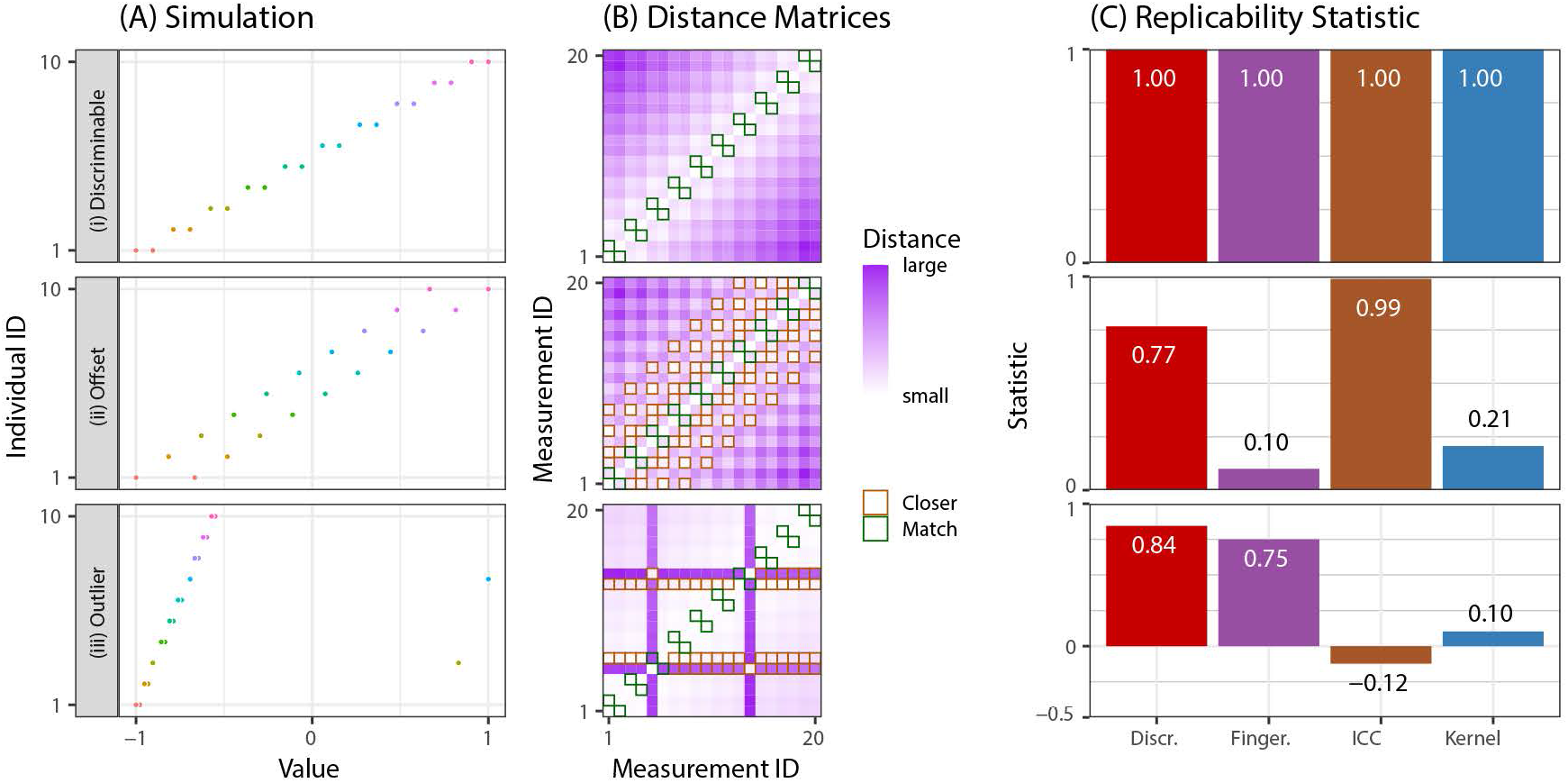
Discr provides a valid discriminability statistic. Three simulations with characteristic notions of discriminability are constructed with *n* = 10 items each with *s* = 2 measurements. **(A)** The 20 samples, where color indicates the individual associated with a single measurement. **(B)** The distance matrices between pairs of measurements. Samples are organized by item. For each row (measurement), green boxes indicate measurements of the same item, and an orange box indicates a measurement from a different item that is more similar to the measurement than the corresponding measurement from the same item. **(C)** Comparison of four replicability statistics in each simulation. Row *(i)*: Each item is most similar to a repeated measurement from the same item. All discriminability statistics are high. Row *(ii)*: Measurements from the same item are more similar than measurements from different individuals on average, but each item has a measurement from a different item in between. `ICC` is essentially unchanged from *(i)* despite the fact that observations from the same individual are less similar than they were in *(i)*, and both `Fingerprint` and `Kernel` are reduced by about an order of magnitude relative to simulation *(i)*. Row *(iii)*: Two of the ten individuals have an “outlier” measurement, and the simulation is otherwise identical to *(i)*. `ICC` is negative, and `Kernel` provides a small statistic. `Discr` is the only statistic that is robust and valid across all of these simulated examples.

#### Definition 1 Inter-Item Discriminability

Assuming we have n items, where each item has s_i_ measurements, we obtain 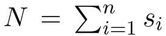 total measurements. For simplicity, assume s_i_ = 2 for the definition below, and that there are no ties. Given that, `Discr` can be computed as follows (for a more formal and general definition and pseudocode, please see Supporting Information S2):

1. Compute the distance between all pairs of samples (resulting in an N × N matrix), Figure 1(B). While any measure of distance is permissible, for the purposes of this manuscript, we perform all our experiments using the Euclidean distance.
2. Identify replicated measurements of the same individual (green boxes). The number of green boxes is g = *n* × 2.
3. For each measurement, identify measurements that are more similar to it than the other measurement of the same item, i.e., measurements whose magnitude is smaller than that in the green box (orange boxes). Let f be the number of orange boxes.
4. Discriminability is defined as fraction of times across-item measurements are smaller than within-item measurements: 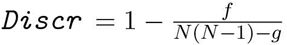.

A high `Discr` indicates that within-item measurements tend to be more similar to one another than across-item measurements. See [39] for a theoretical analysis of `Discr` as compared to these and other data replicability statistics. For brevity, we use the term “discriminability” to refer to inter-item discriminability hereafter.

### 2.2 Testing for discriminability

Letting *R* denote the replicability of a dataset with *n* items and *s* measurements per item, and *R*_0_ denote the replicability of the same size dataset with zero item specific information, test for replicability is

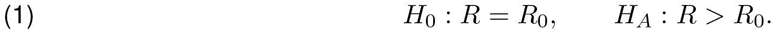

One can use any ‘data replicability’ statistic for *R* and *R*_0_ [39]. We devised a permutation test to obtain a distribution of the test statistic under the null, and a corresponding p-value. To evaluate the different procedures, we compute the power of each test, that is, the probability of correctly rejecting the null when it is false (which is one minus type II error; see Supporting Information S5.1 for details).

### 2.3 Testing for better discriminability

Letting *R*^(1)^ be the replicability of one dataset or approach, and *R*^(2)^ be the replicability of the second, we have the following comparison hypothesis for replicability:

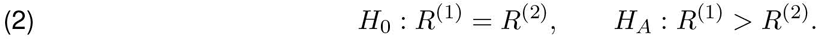

Again, we devised a permutation test to obtain the distribution of the test statistic under the null, and p-values (see Supporting Information S5.2 for details).

### 2.4 Simulation settings

To develop insight into the performance of `Discr`, we consider several different simulation settings (see Supporting Information S4 for details). Each setting includes between 2 and 20 items, with 128 total measurements, in two dimensions:

1. **Gaussian** Sixteen items are each distributed according to a spherically symmetric Gaussian, therefore respecting the assumptions that motivate intraclass correlations.
2. **Cross** Two items have Gaussian distributions with the same mean and different diagonal covariance matrices.
3. **Ball/Circle** One item is distributed in the unit ball, the other on the unit circle; Gaussian noise is added to both.
4. **XOR** Each of two items is a mixture of two spherically symmetric Gaussians, but means are organized in an XOR fashion; that is, the means of the first item are (0, 1) and (1, 0), whereas the means of the second are (0, 0) and (1, 1). The implication is that many measurements from a given item are further away than any measurement of the other item.
5. **No Signal** Both items have the same Gaussian distribution.

## 3 Results

### 3.1 Theoretical properties of Discriminability

Under reasonably general assumptions, if within-item variability increases, predictive accuracy will decrease. Therefore, a statistic that is sensitive to within-item variance is desirable for optimal experimental design, regardless of the distribution of the data. [40] introduces a univariate parametric framework in which predictive accuracy can be lower-bounded by a decreasing function of `ICC`; as a direct consequence, a strategy with a higher `ICC` will, on average, have higher predictive performance on subsequent inference tasks. Unfortunately, this valuable theoretical result is limited in its applicability, as it is restricted to univariate data, whereas big data analysis strategies often produce multivariate data. We therefore prove the following generalization of this theorem:

#### Theorem 1.

Under the multivariate mixture model with the first two moments bounded above, plus additive noise setting, or a sufficient generalization thereof, Discr provides a lower bound on the predictive accuracy of a subsequent classification task. Consequently, a strategy with a higher Discr provably provides a higher bound on predictive accuracy than a strategy with a lower `Discr`.

See Supporting Information S3 for proof. Correspondingly, this property motivates optimizing experiments to obtain higher Discr.

### 3.2 Properties of various replicability statistics

In Fig 1, we highlight the properties of different statistics across a range of basic one-dimensional simulations, all of which display a characteristic notion of replicability: samples of the same item tend to be more similar to one another than samples from different items. In three different univariate simulations we observe two samples from ten items (Figure 1A), and the construct in which replicability statistics will be evaluated:

i. **Discriminable** has each item’s samples closer to each other than any other items. The replicability statistic should attain a large value to reflect the high within-item similarity compared to the between-item similarity.
ii. **Offset** shifts the second measurement a bit, so that it is further from the first measurement than another item. Replicability statistic should still be high, but lower than the offset simulation.
iii. **Outlier** is the same as **discriminable** but includes two items with an outlier measurement. This is another highly reliable setting, so we hope outliers do not significantly reduce the replicability score.

We compare `Discr` to intraclass correlation coefficient (`ICC`), fingerprinting index (`Fingerprint`) [37], and k-sample kernel testing (`Kernel`) [41] (see Supporting Information S1 for details). `ICC` provides no ability for differentiating between *discriminable* and *offset* simulation, despite the fact that the data in *discriminable* is more replicable than *offset*. While this property may be useful in some contexts, a lack of sensitivity to the offset renders users unable to discern which strategy has a higher test-retest reliability. Moreover, `ICC` is uninterpretable in the case of even a very small number of outliers, where `ICC` is negative. On the other hand, `Fingerprint` suffers from the limitation that if the nearest measurement is anything but a measurement of the same item, it will be at or near zero, as shown in *offset*. `Kernel` also performs poorly in *offset* and in the presence of *outliers*. In contrast, across all simulations, `Discr` shows reasonable construct validity under the given constructs: the statistic is high across all simulations, and highest when repeated measurements of the same item are more similar than measurements from any of the other items.

### 3.3 The power of replicability statistics in multivariate experimental design

We evaluate `Discr`, `PICC` (which applies `ICC` to the top principal component of the data), `I2C2`, `Fingerprint`, and `Kernel` on five two-dimensional simulation settings (see Supporting Information S4 for details). Fig 2A shows a two-dimensional scatterplot of each setting, and Fig 2B shows the Euclidean distance matrix between samples, ordered by item.

**Fig 2.**
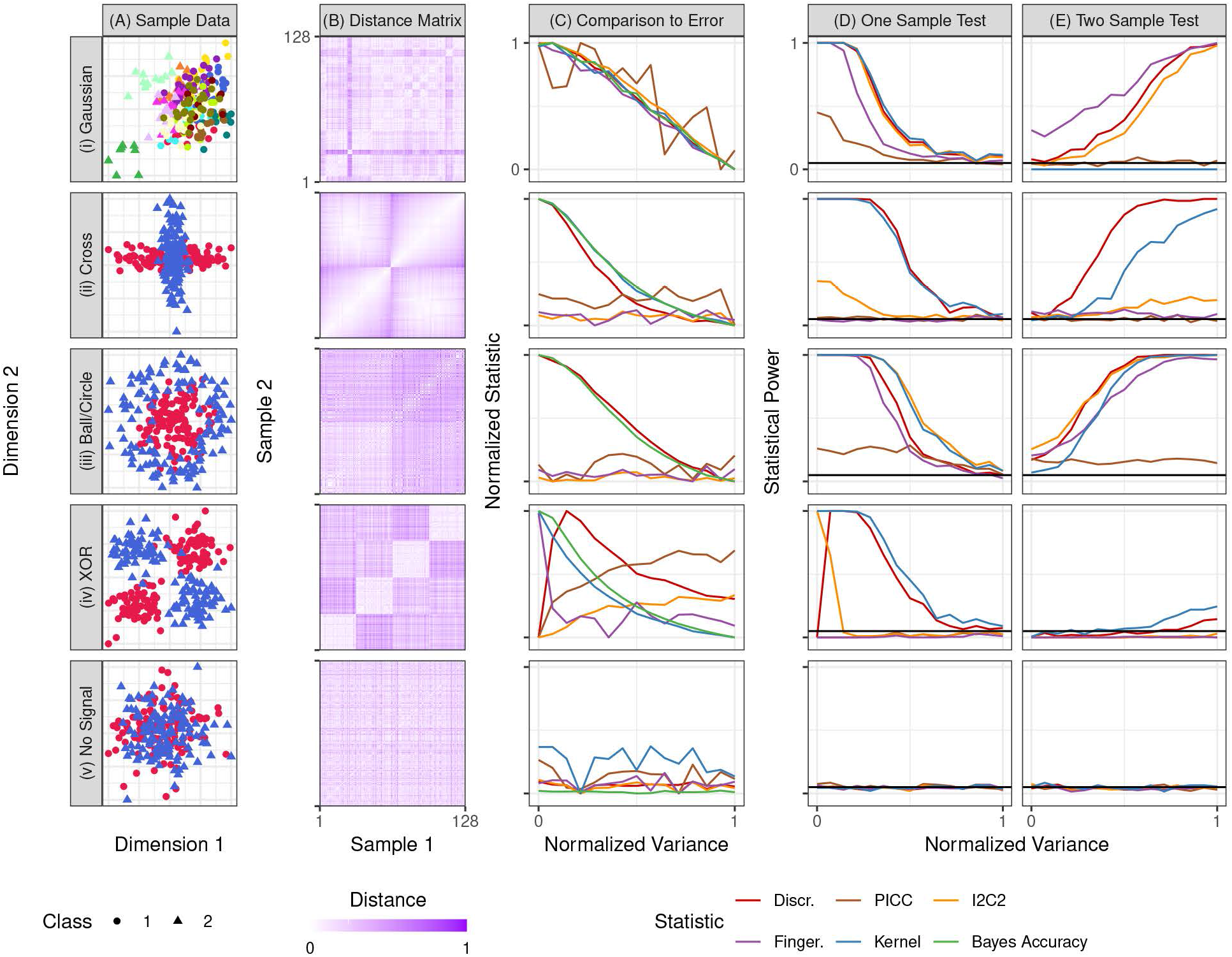
Multivariate simulations demonstrate the value of optimizing replicability for experimental design. All simulations are two-dimensional, with 128 samples, with 500 iterations per setting (see Supporting Information S4 for details). **(A)** For each setting, class label is indicated by shape, and color indicates item identity. **(B)** Euclidean distance matrix between samples within each simulation setting. Samples are organized by item. Simulation settings in which items are discriminable tend to have a block structure where samples from the same item are relatively similar to one another. **(C)** Replicability statistic versus variance. Here, we can compute the Bayes accuracy (the best one could perform to predict class label) as a function of variance. `Discr` and `Kernel` are mostly monotonic relative to within-item variance across all settings, suggesting that one can predict improved performance via improved `Discr`. **(D)** Test of whether data are discriminable. `Discr` typically achieves high power among the alternative statistics in all cases. **(E)** Comparison test of which approach is more discriminable. `Discr` is the only statistic which achieves high power in all settings in which any statistic was able to achieve high power.

#### Discriminability empirically predicts performance on subsequent inference tasks

Fig 2C shows the impact of increasing within-item variance on the different simulation settings. The purpose of these simulations is to assess the degree to which `Discr` or the other replicability statistics correspond to downstream predictive accuracy, both under a multivariate Gaussian assumption, and more generally. For the top four simulations, increasing variance decreases predictive accuracy (green line). As desired, `Discr` also decreases nearly perfectly monotonically with decreasing variances. However, only in the first setting, where each item has a spherically symmetric Gaussian distribution, do `I2C2`, `PICC`, and `Fingerprint` drop proportionally. Even in the second (Gaussian) setting, `I2C2`, `PICC`, and `Fingerprint` are effectively uninformative about the within-item variance. And in the third and fourth (non-Gaussian) settings, they are similarly useless. In the fifth simulation they are all at chance levels, as they should be, because there is no information about class in the data. This suggests that of these statistics, only `Discr` and `Kernel` can serve as satisfactory surrogates for predictive accuracy under these quite simple settings.

#### A test to determine replicability

A prerequisite for making item-specific predictions is that items are different from one another in predictable ways, that is, are discriminable. If not, the same assay applied to the same individual on multiple trials could yield unacceptably highly variable results. Thus, prior to embarking on a machine learning search for predictive accuracy, one can simply test whether the data are discriminable at all. If not, predictive accuracy will be hopeless.

Fig 2D shows that `Discr` achieves high power among all competing approaches in all settings and variances. This result demonstrates that despite the fact that `Discr` does not rely on Gaussian assumptions, it still performs nearly as well or better than parametric methods when the data satisfy these assumptions (row (i)). In row (ii) cross, only `Discr` and `Kernel` correctly identify that items differ from one another, despite the fact that the data are Gaussian, though they are not spherically symmetric gaussians. In both rows (iii) ball/disc and (iv) XOR, most statistics perform well despite the non-Gaussianity of the data. And when there is no signal, all tests are valid, achieving power less than or equal to the critical value. Non-parametric `Discr` therefore has the power of parametric approaches for data at which those assumptions are appropriate, and much higher power for other data. `Kernel` performs comparably to `Discr` in these settings.

#### A test to compare reliabilities

Given two experimental designs—which can differ either by acquisition and/or analysis details—are the measurements produced by one method more discriminable than the other? Fig 2D shows `Discr` typically achieves the highest power among all statistics considered. Specifically, only `Fingerprint` achieves higher power in the Gaussian setting, but it achieves almost no power in the cross setting. `Kernel` achieves comparably lower power for most settings and no power for the Gaussian, as does `PICC`. `I2C2` achieves similar power to `Discr` only for the Gaussian and ball/disc setting. All tests are valid in that they achieve a power approximately equal to or below the critical value when there is no signal. Note that these comparisons are not the typical “k-sample comparisons” with many theoretical results, rather, they are comparing across multiple disparate k-sample settings. Thus, in general, there is a lack of theoretical guarantees for this setting. Nonetheless, the fact that `Discr` achieves nearly equal or higher power than the statistics that build upon Gaussian methods, even under Gaussian assumptions, suggests that `Discr` will be a superior metric for optimal experimental design in real data.

### 3.4 Optimizing experimental design via maximizing replicability in human brain imaging data

#### Human brain imaging data acquisition and analysis

Consortium for Reliability and Reproducibility (CoRR) [42] has generated functional, anatomical, and diffusion magnetic resonance imaging (dMRI) scans from > 1,600 participants, often with multiple measurements, collected through 28 different datasets (22 of which have both age and sex annotation) spanning over 20 sites. Each of the sites use different scanners, technicians, scanning protocols, and retest follow up procedures, thereby representing a wide variety of different acquisition settings with which one can test different analysis pipelines. Supporting Information S6.3 provides protocol metadata associated with each individual dataset. Fig 3A shows the six stage sequence of analysis steps for converting the raw fMRI data into networks or connectomes, that is, estimates of the strength of connections between all pairs of brain regions. At each stage of the pipeline, we consider several different “standard” approaches, that is, approaches that have previously been proposed in the literature, typically with hundreds or thousands of citations [43]. Moreover, they have all been collected into an analysis engine, called Configurable Pipeline for the Analysis of Connectomes (C-PAC) [44]. In total, for the six stages together, we consider 2 × 2 × 2 × 2 × 4 × 3 = 192 different analysis pipelines. Because each stage is nonlinear, it is possible that the best sequence of choices is not equivalent to the best choices on their own. For this reason, publications that evaluate a given stage using any metric, could result in misleading conclusions if one is searching for the best sequence of steps [45]. The dMRI connectomes were acquired via 48 analysis pipelines using the Neurodata MRI Graphs (`ndmg`) pipeline [46]. Supporting Information S6 provides specific details for both fMRI and dMRI analysis, as well as the options attempted.

**Fig 3.**
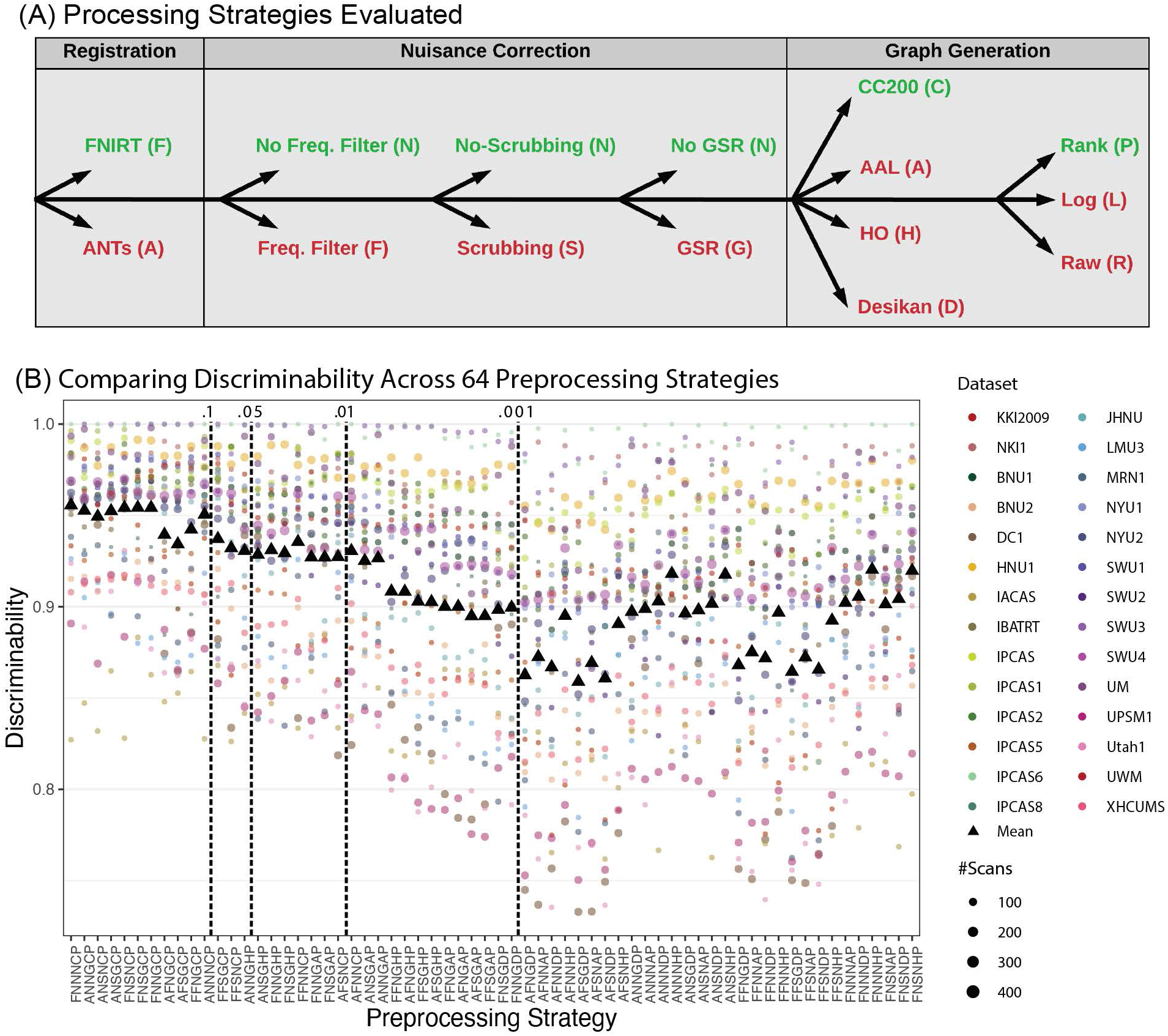
Different analysis strategies yield widely disparate stabilities. **(A)** Illustration of analysis options for the 192 fMRI pipelines under consideration (described in Supporting Information S6.1). The sequence of options corresponding to the best performing pipeline overall are in green. **(B)** `Discr` of fMRI Connectomes analyzed using 64 different pipelines. Functional correlation matrices are estimated from 28 multi-session studies from the CoRR dataset using each pipeline. The analysis strategy codes are assigned sequentially according to the abbreviations listed for each step in **(A)**. The mean `Discr` per pipeline is a weighted sum of its stabilities across datasets. Each pipeline is compared to the optimal pipeline with the highest mean `Discr`, `FNNNCP`, using the above comparison hypothesis test. The remaining strategies are arranged according to *p*-value, indicated in the top row.

#### Different analysis strategies yield widely disparate stabilities

The analysis strategy has a large impact on the `Discr` of the resulting fMRI connectomes (Fig 3B). Each column shows one of 64 different analysis strategies, ordered by how significantly different they are from the pipeline with highest `Discr` (averaged over all datasets, tested using the above comparison test). Interestingly, pipelines with worse average `Discr` also tend to have higher variance across datasets. The best pipeline, FNNNCP, uses FSL registration, no frequency filtering, no scrubbing, no global signal regression, CC200 parcellation, and converts edges weights to ranks. While all strategies across all datasets with multiple participants are significantly discriminable at *α* = 0.05 (`Discr` goodness of fit test), the majority of the strategies (51/64 ≈ 80%) show significantly worse Discr than the optimal strategy at *α* = 0.05 (`Discr` comparison test).

#### Discriminability identifies which acquisition and analysis decisions are most important for improving performance

While the above analysis provides evidence for which *sequence* of analysis steps is best, it does not provide information about which choices individually have the largest impact on overall `Discr`. To do so, it is inadequate to simply fix a pipeline and only swap out algorithms for a single stage, as such an analysis will only provide information about that fixed pipeline. Therefore, we evaluate each choice in the context of all 192 considered pipelines in Fig 4A. The pipeline constructed by identifying the best option for each analysis stage is `FNNGCP` (Figure 4A). Although it is not exactly the same as the pipeline with highest `Discr` (`FNNNCP`), it is also not much worse (`Discr` 2-sample test, p-value ≈ 0.14). Moreover, except for scrubbing, each stage has a significant impact on `Discr` after correction for multiple hypotheses (Wilcoxon signed-rank statistic, *p*-values all < 0.001).

**Fig 4.**
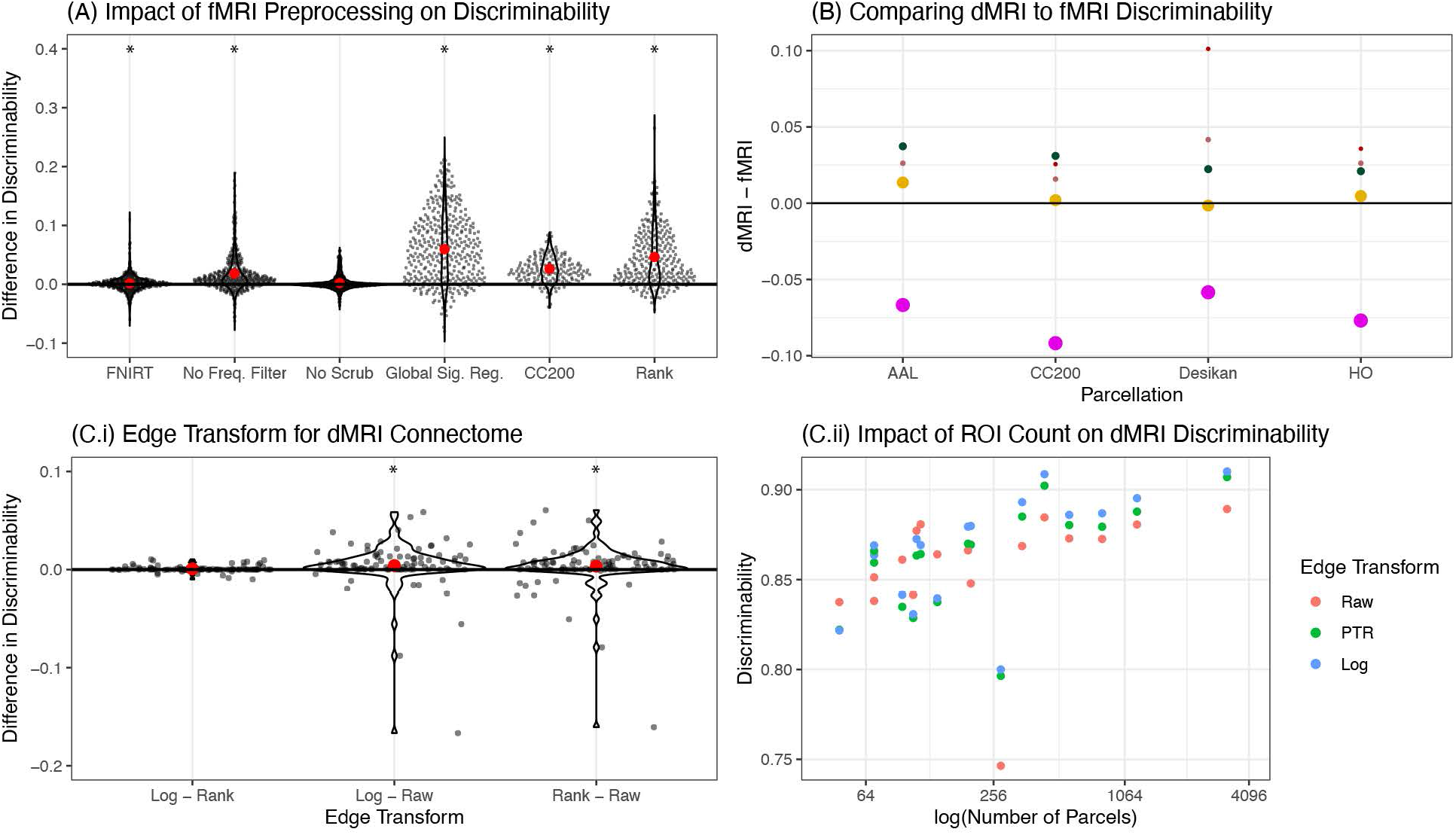
Parsing the relative impact on Discr of various acquisition and analytic choices. **(A)** The pipelines are aggregated for a particular analysis step, with pairwise comparisons with the remaining analysis options held fixed. The beeswarm plot shows the difference between the overall best performing option and the second best option for each stage (mean in red) with other options held equal; the *x*-axis label indicates the best performing strategy. The best strategies are `FNIRT`, no frequency filtering, no scrubbing, global signal regression, the CC200 parcellation, and ranks edge transformation. A Wilcoxon signed-rank test is used to determine whether the mean for the best strategy exceeds the second best strategy: a * indicates that the *p*-value is at most 0.001 after Bonferroni correction. Of the best options, only no scrubbing is *not* significantly better than alternative strategies. Note that the options that perform marginally the best are not significantly different than the best performing strategy overall, as shown in Fig 3. **(B)** A comparison of the stabilities for the 4 datasets with both fMRI and dMRI connectomes. dMRI connectomes tend to be more discriminable, in 14 of 20 total comparisons. Color and point size correspond to the study and number of scans, respectively (see Fig 3B). **(C.i)** Comparing raw edge weights (Raw), ranking (Rank), and log-transforming the edge-weights (Log) for the diffusion connectomes, the Log and Rank transformed edge-weights tend to show higher Discr than Raw. **(C.ii)** As the number of ROIs increases, the `Discr` tends to increase.

Another choice is whether to estimate connectomes using functional or diffusion MRI (Figure 4B). Whereas both data acquisition strategies have known problems [47], the `Discr` of the two experimental modalities has not been directly compared. Using four datasets from CoRR that acquired both fMRI and dMRI on the same subjects, and have quite similar demographic profiles, we tested whether fMRI or dMRI derived connectomes were more discriminable. The pipelines being considered were the best-performing fMRI pre-processing pipeline (`FNNNCP`) against the dMRI pipeline with the CC200 parcellation. For three of the four datasets, dMRI connectomes were more discriminable. This is not particularly surprising, given the susceptibility of fMRI data to changes in state rather than trait (e.g., amount of caffeine prior to scan [44]).

The above results motivate investigating which aspects of the dMRI analysis strategy were most effective. We focus on two criteria: how to scale the weights of connections, and how many regions of interest (ROIs) to use. For scaling the weights of the connections, we consider three possible criteria: using the raw edge-weights (“Raw”), taking the log of the edge-weights (“Log”), and ranking the nonzero edge weights in sequentially increasing order (“Rank”). Fig 4C.i shows that both rank and log transform significantly exceed raw edge weights (Wilcoxon signed-rank statistic, sample size= 60, p-values all < 0.001). Fig 4C.ii shows that parcellations with larger numbers of ROIs tend to have higher `Discr`. Unfortunately, most parcellations with semantic labels (e.g., visual cortex) have hundreds not thousands of parcels. This result therefore motivates the development of more refined semantic labels.

#### Optimizing Discriminability improves downstream inference performance

We next examined the relationship between the `Discr` of each pipeline, and the amount of information it preserves about two properties of interest: sex and age. Based on the simulations above, we expect that analysis pipelines with higher `Discr` will yield connectomes with more information about covariates. Indeed, Fig 5 shows that, for virtually every single dataset including sex and age annotation (22 in total), a pipeline with higher `Discr` tends to preserve more information about both covariates. The amount of information is quantified by the effect size of the distance correlation `DCorr` (which is exactly equivalent to `Kernel` [36, 48]), a statistic that quantifies the magnitude of association for both linear and nonlinear dependence structures. In contrast, if one were to use either `Kernel` or `I2C2` to select the optimal pipeline, for many datasets, subsequent predictive performance would degrade. `Fingerprint` performs similarly to `Discr`, while `PICC` provides a slight decrease in performance on this dataset. These results are highly statistically significant: the slopes of effect size versus `Discr` and `Fingerprint` across datasets are significantly positive for both age and sex in 82 and 95 percent of all studies, respectively (robust *Z*-test, *α* = 0.05). `Kernel` performs poorly, basically always, because *k*-sample tests are designed to perform well with many samples from a small number of different populations, and questions of replicability across repeated measurements have a few samples across many different populations.

**Fig 5.**
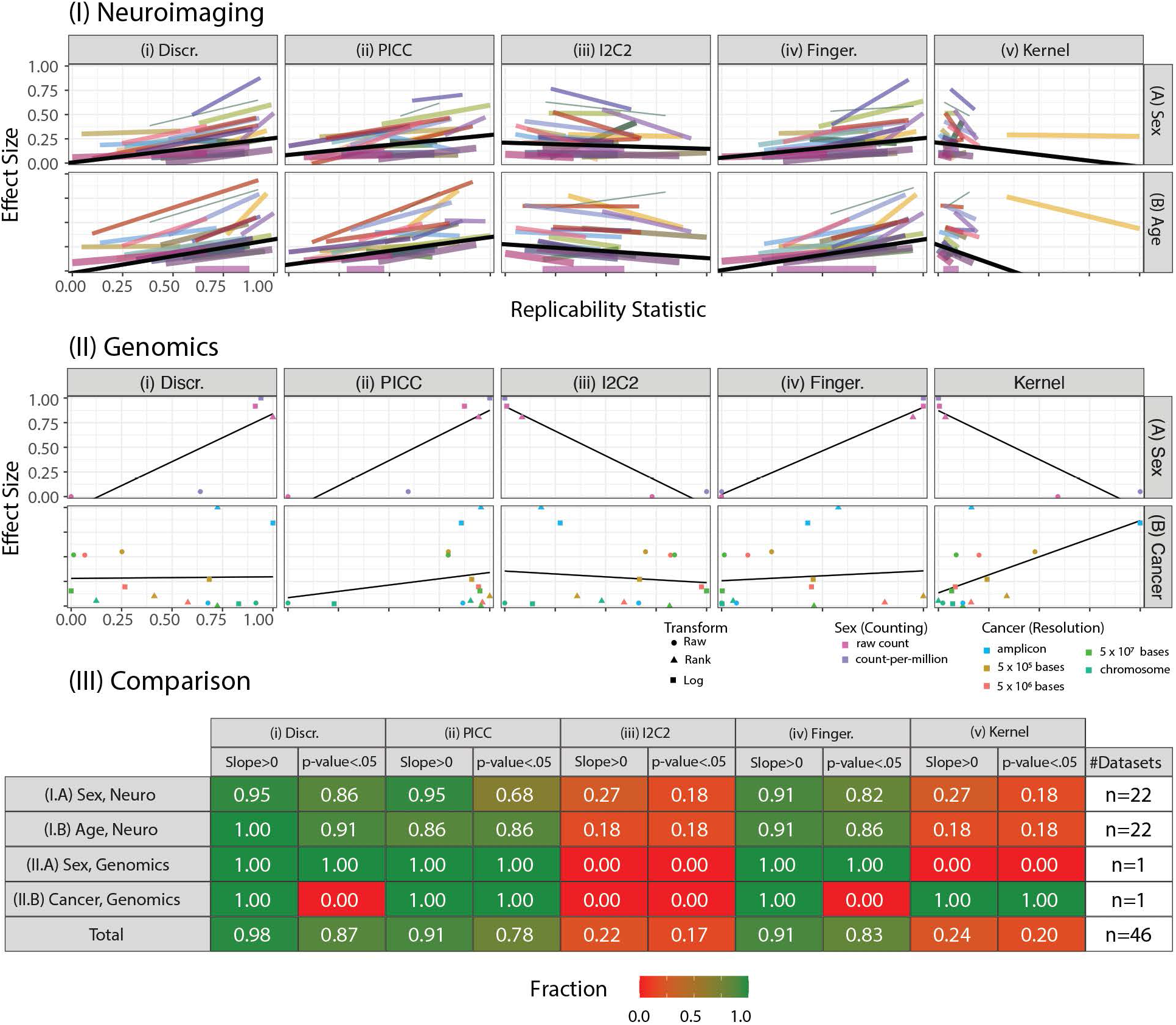
Optimizing Discr improves downstream inference performance. Using the connectomes from the 64 pipelines with raw edge-weights, we examine the relationship between connectomes vs sex and age. The columns evaluate difference approaches for computing pipeline effectiveness, including **(i)** `Discr`, **(ii)** `PICC`, **(iii)** Average Fingerprint Index `Fingerprint`, **(iv)** `I2C2`, and **(v)** `Kernel`. Each panel shows reference pipeline replicability estimate (*x-axis*) versus effect size of the association between the data and the sex, age, or cancer status of the individual as measured by `DCorr` (*y-axis*). Both the *x* and *y* axes are normalized by the minimum and maximum statistic. These data are summarized by a single line per study, which is the regression of the normalized effect size onto the normalized replicability estimate as quantified by the indicated reference statistic. **(I)** The results for the neuroimaging data, as described in Section 3.4. Color and line width correspond to the study and number of scans, respectively (see Fig 3B). The solid black line is the weighted mean over all studies. `Discr` is the only statistic in which *nearly all* slopes are positive. Moreover, the corrected *p*-value [51, 52] is significant across most datasets for both covariates (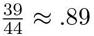 *p*4-values < .001). This indicates that pipelines with higher `Discr` correspond to larger effect sizes for the covariate of interest, and that this relationship is stronger for `Discr` than other statistics. A similar experiment is performed on two genomics datasets, measuring the effects due to sex and whether an individual has cancer. **(III)** indicates the fraction of datasets with positive slopes and with significantly positive slopes, ranging from 0 (“None”, red) to 1 (“All”, green), at both the task and aggregate level. `Discr` is the statistic where the most datasets have positive slopes, and the statistic where the most datasets have significantly positive slopes, across the neuroimaging and genomics datasets considered. Supporting Information S6.2 details the methodologies employed.

### 3.5 Reliability of genomics data

The first genomics study aimed to explore variation in gene expression across human induced pluripotent stem cell (hiPSC) lines with between one and seven replicates [49]. This data includes RNAseq data from 101 healthy individuals, comprising 38 males and 63 females. Expression was interrogated across donors by studying up to seven replicated iPSC lines from each donor, yielding bulk RNAseq data from a total of 317 individual hiPSC lines. While the pipeline includes many steps, we focus here for simplicity on (1) counting, and (2) normalizing. The two counting approaches we study are the raw hiPSC lines and the count-per-million (CPM). Given counts, we consider three different normalization options: Raw, Rank, and Log-transformed (as described above). The task of interest was to identify the sex of the individual.

The second genomics study [50] includes 331 individuals, consisting of 135 patients with non-metastatic cancer and 196 healthy controls, each with eight DNA samples. The study leverages a PCR-based assay called Repetitive element aneuploidy sequencing system to analyze 750,000 amplicons distributed throughout the genome to investigate the presence of aneuploidy (abnormal chromosome counts) in samples from cancer patients (see Supporting Information S6.1 for more details). The possible processing strategies include using the raw amplicons or the amplicons downsampled by a factor of 5 × 10^5^ bases, 5 × 10^6^ bases, 5 × 10^7^ bases, or to the individual chromosome level (the *resolution* of the data), followed by normalizing through the previously described approaches (Raw, Rank, Log-transformed) yielding 5 × 3 = 15 possible strategies in total. The task of interest was to identify whether the sample was collected from a cancer patient or a healthy control.

Across both tasks, slope for discriminability is positive, and for the first task, the slope is significantly bigger than zero (robust *Z*-test, *p*-value = .001, *α* = .05). `Fingerprint` and `Kernel` are similarly only informative for one of the two genomics studies. For `PICC`, in both datasets the slope is positive and the effect is significant. `I2C2` does not provide value for subsequent inference.

## 4 Discussion

We propose the use of the `Discr` statistic as a simple and intuitive measure for experimental design featuring multiple measurements. Numerous efforts have established the value of *quantifying* repeatability and replicability (or discriminability) using parametric measures such as `ICC` and `I2C2`. However, they have not been used to optimize replicability—that is, they are only used post-hoc to determine replicability, not used as criteria for searching over the design space—nor have non-parametric multivariate generalizations of these statistics been available. We derive goodness of fit and comparison (equality) tests for `Discr`, and demonstrate via theory and simulation that `Discr` provides numerous advantages over existing techniques across a range of simulated settings. Our neuroimaging and genomics use-cases exemplify the utility of these features of the `Discr` framework for optimal experimental design.

An important consideration is that quantifying reliability and replicability with multiple measurements may seem like a limitation for many fields, in which the end derivative typically used for inference may be just a single sample for each item measured. However, a single measurement may often consist of many sub-measurements for a single individual, each of which are combined to produce the single derivative work. For example in brain imaging, a functional Magnetic Resonance Imaging (fMRI) scan consists of tens to thousands of scans of the brain at numerous time points. In this case, the image can be broken into identical-width time windows to coerce a dataset in which discriminability can be investigated. In another example taken directly from the cancer genomics experiment below, a genomics count table was produced from eight independent experiments, each of which yielded a single count table. The last step of their pre-processing procedure was to aggregate to produce the single summary derivative that the experimenters traditionally considered a single measurement. In each case, the typical “measurement” unit can really be thought of as an aggregate of multiple smaller measurement units, and a researcher can leverage these smaller measurements as a surrogate for multiple measurements. In the neuroimaging example, the fMRI scan can be segmented into identical-width sub-scans with each treated as a single measurement, and in the genomics example, the independent experiments can each be used as a single measurement.

`Discr` provides a number of connections with related statistical algorithms worth further consideration. `Discr` is related to energy statistics [53], in which the statistic is a function of distances between observations [33]. Energy statistics provide approaches for goodness-of-fit (one-sample) and equality testing (two-sample), and multi-sample testing [54]. However, we note an important distinction: a goodness of fit test for discriminability can be thought of as a *K*-sample test in the classical literature, and a comparison of discriminabilities is analogous to a comparison of *K*-sample tests. Further, similar to `Discr`, energy statistics make relatively few assumptions. However, energy statistics requires a large number of measurements per item, which is often unsuitable for biological data where we frequently have only a small number of repeated measurements. `Discr` is most closely related to multiscale generalized correlation (`MGC`) [36, 48], which combines energy statistics with nearest neighbors, as does `Discr`. Like many energy-based statistics, `Discr` relies upon the construction of a distance matrix. As such, `Discr` generalizes readily to high-dimensional data, and many packages accelerate distance computation in high-dimensionals [55].

### Limitations

While `Discr` provides experimental design guidance for big data, other considerations may play a role in a final determination of the practical utility of an experimental design. For example, the connectomes analyzed here are *resting-state*, as opposed to *task-based* fMRI connectomes. Recent literature suggests that the global signal in a rs-fMRI scan may be correlated heavily with signals of interest for task-based approaches [56, 57], and therefore removal may be inadvisable. Thus, while `Discr` is an effective tool for experimental design, knowledge of the techniques in conjunction with the constructs under which successive inference will be performed remains essential. Further, in this study, we only consider the Euclidean distance, which may not be appropriate for all datasets of interest. For example, if the measurements live in a manifold (such as images, text, speech, and networks), one may be interested in dissimilarity or similarity functions other than Euclidean distance. To this end, Discr readily generalizes to alternative comparison functions, and will produce an informative result as long as the choice of comparison function is appropriate for the measurements.

It is important to emphasize that `Discr`, as well the related statistics, are neither necessary, nor sufficient, for a measurement to be practically useful. For example, categorical covariates, such as sex, are often meaningful in an analysis, but not discriminable. Human fingerprints are discriminable, but typically not biologically useful. In this sense, while discriminability provides a valuable link between test-retest reliability and criterion validity for multivariate data, one must be careful to consider other notions of validity prior to the selection of a measurement. In addition, none of the statistics studied here are immune to sample characteristics, thus interpreting results across studies deserves careful scrutiny. For example, having a sample with variable ages will increase the inter-subject dissimilarity of any metric dependent on age (such as the connectome). Additionally, discriminability can be decomposed into within and between-class discriminabilities, so that class-specific effects may be examined in isolation, as described in Supporting Information S7. Future work could explore how these two quantities may be incorporated into the experimental design procedure.

Moreover, if multiple strategies are saturated at a perfect discriminability (`Discr` = 1), it does not provide an informative way to differentiate between these strategies. One could trivially augment the discriminability procedure to compare within-item distances to a scaled and/or shifted transformation of between-item distances, thereby rendering perfect discriminability arbitrarily difficult. With these caveats in mind, `Discr` remains a key experimental design consideration across a wide variety of settings.

### Conclusion

The use-cases provided herein serve to illustrate how `Discr` can be used to facilitate experimental design, and mitigate replicability issues. We envision that `Discr` will find substantial applicability across disciplines and sectors beyond brain imaging and genomics, such pharmaceutical research. To this end, we provide open-source implementations of `Discr` for both `Python` and `R` [58, 59]. Code for reproducing all the figures in this manuscript is available at https://neurodata.io/mgc.

## Acknowledgements

The authors would like to thank Iris Van Rooij and the Neurodata team for their valuable feedback on this manuscript.

---

Supporting Information Legend.

Supporting Information Legend
| Section | Description |
| --- | --- |
| S1 | Background information on repeatability statistics. |
| S2 | Population and sample discriminability. |
| S3 | Theoretical bound for downstream inference. |
| S4 | Simulation settings. |
| S5 | Hypothesis testing. |
| S6 | Data descriptions and details for real data analysis. |
| S7 | Extensions of discriminability. |

## Supporting Information 1

### S1 Data Repeatability Statistics

#### Intraclass Correlation Coefficient

The intraclass correlation coefficient (`ICC`) is a commonly used data replicability statistic [1]. The absolute agreement `ICC`, or `ICC(1,1)`, is the fraction of the total variability that is across-item variability, that is, `ICC` is defined as the across-item variability divided by the within-item plus across-item variability. `ICC` has several limitations. First, it is univariate, meaning if the data are multidimensional, they must first be represented by univariate statistics, thereby discarding multivariate information. This potentially makes `ICC` unsuitable when an informative univariate summary measure is unavailable or unknown, which is frequently the case in the high dimensional data that is the focus of this manuscript. Second, `ICC` is based on a Gaussian assumption characterizing the data. Thus, any deviations from this assumption may render the interpretation of the magnitude of `ICC` questionable, because non-Gaussian measurements that are highly replicable could potentially yield quite low `ICC` [2–4]. Third, the Intraclass correlation coefficient is highly sensitive to the design of the study [4, 5]; care must be taken to ensure that the form of ICC chosen accurately reflects the design of the study of interest. Further, `ICC` is substantially impacted by the presence of outliers in measurements [6]. Finally, there are numerous definitions of estimates of `ICC`[1] designed for different experimental setups, and researchers regularly use (and misuse) the different estimators in generic contexts [4, 7]. In practice, it is unclear the extent to which the use of inappropriate estimators of `ICC` is impactful [8].

Numerous multivariate generalizations of the `ICC` attempt to overcome the requirement of `ICC` to operate on univariate data. The Image Intra-Class Correlation (`I2C2`) was introduced to mitigate `ICC`’s univariate limitation [9]. Specifically, `I2C2` operates on covariances matrices, rather than variances. To obtain a univariate summary of replicability, `I2C2` operates on the trace of the covariance matrices, one of several possible strategies, similar to most multivariate analysis of variance procedures [10]. Thus, while overcoming one limitation of `ICC`, `I2C2` still heavily leverages Gaussian assumptions of the data to justify its validity. [11] highlight a number of limitations with using estimates of covariance in the context of assessing multivariate replicability. Chiefly, sampling variance of covariance components in the high dimensionality; low-sample-size (HDLSS) regime is problematic, which is an characteristic of increasing prevalence in biological data.

#### Fingerprinting Index

The fingerprinting index [12, 13] provides a metric for quantifying individual connectivity profiles in resting-state MRI (fMRI). Specifically, the fingerprinting index operates on the pairwise correlation of the vectorized connectivity matrices. A high fingerprinting index corresponds to the connectivity matrices being most strongly correlated within-subject versus between-subject. An important clarification for fingerprinting is that the connectivity matrices must be more strongly correlated than *any other measurement* within a particular scanning session, otherwise the fingerprinting index will be 0, as the fingerprinting index uses only the nearest-neighbor associated with a given item. Unlike the other strategies employed in this manuscript, the fingerprinting index produces a statistic for each possible ordering of 2 measurement sessions, that is, if each item is measured *s* times, fingerprinting produces *s*(*s* − 1) statistics. To enable fingerprinting for assessing the effectiveness of a strategy, we instead averaged across all *s*(*s* − 1) statistics, which will henceforth be referred to as `Fingerprinting`.

#### Kendall’s Coefficient of Concordance

Kendall’s Coefficient of Concordance, or Kendall’s *W* , is a univariate non-parametric statistic for assessing the extent to which multiple measurements of the same item agree. Like inter-item discriminability and the fingerprinting index, estimates of Kendall’s *W* operate on the ranks of data. Specifically, Kendall’s *W* computes the total rank of all measurements associated with a single item, and compares an item’s total rank to the average value of the total rank. An important consideration is that Kendall’s *W* operates directly on the measurements themselves, rather than on scalar summary measures of the relationships amongst the measurements. As such, Kendall’s *W* cannot be applied directly to data that is inherently multivariate using traditional methods of ranking. For this reason, we do not formally evaluate Kendall’s *W* within the context of this manuscript.

#### Kernel Methods

Maximum mean discrepancy (`MMD`) [14] provides a non-parametric framework for comparing whether two samples are drawn from the same distribution. `MMD` subverts Gaussian assumptions by embedding the points in a reproducing kernel Hilbert Space (RKHS), and looking for functions over the unit ball in the RKHS which maximize the difference in the means of the embedded points. In the two-item regime, `MMD` can be shown to be equivalent to the Hilbert-Schmidt Independence Criterion (`HSIC`) [15–17], which provides a natural generalization of `MMD` when the number of classes exceeds two. To date, to our knowledge, there does not exist a k-sample variant of `MMD`.

Distance Components (`DISCO`) [18] extends the classical Analysis of Variance (`ANOVA`) framework to cases where the distributions are not necessarily Gaussian. In contrast to `ANOVA` which makes simplifying assumptions of normality, `DISCO` operates on the dispersion of the samples based on the Euclidean Distance, comparing the within-class dispersion to the between-class dispersion. `DISCO` produces a consistent test against general alternatives as the number of observations *s* per item goes to infinity. [19] shows a closed form relationship between `Kernel` and other Energy statistics approaches, such as Distance correlation. The result is that using Distance correlation for k-sample testing results in a test statistic that has bias relative to the Kernel statistic, but will yield the same p-value. Further, [19] shows the equivalence between Distance correlation and `HSIC/MMD`. Thus, in this manuscript, we use Kernel to refer to either `DISCO` or `MMD` as appropriate. In all cases, we use the default kernel, which is the Gaussian kernel with the typical bandwidth specification, as implemented in the `kernlab` package [20] (`MMD`) and energy (`DISCO`) package [21]. Note that in many real data scenarios, *s* is small (particularly, most “repeat measurements” datasets have *s* = 2), and the finite-sample performance of `Kernel` on such a small number of repeat trials is not known.

## Supporting Information 2

### S2 Population and Sample Discr

Suppose that ***θ**_i_* ∈ Θ represents a physical property of interest for a particular item *i*. In a biological context, for instance, an item could be a participant in a study, and the property of interest could be the individual’s true brain network, or connectome. We cannot directly observe the physical property, but rather, we must first measure ***θ**_i_* and then “wrangle” it. Call the measurement function, 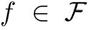 for a family of possible measurement functions 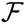 That is, 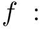 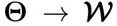. So, measurements of ***θ**_i_* are observed as *f* (***θ**_i_*) = *w_i_*. However, *w_i_* may be a noisy, with measurement artefacts. Alternately, *w_i_* might not be the property of interest, for example, if the property is a network, perhaps *w_i_* is a multivariate time-series, from which we can estimate a network. We therefore have another function, 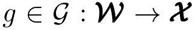, which represents the data wrangling procedure to take the measurement and produce an informative derivative (for instance, confound removal). The family of possible data wrangling procedures to produce the informative derivative is 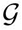. In this fashion, the output of interest is *x_i_* = *g*(*f* (***θ**_i_*)).

The goal of experimental design is to choose an *f* and *g* that yield high-quality and useful inferences, that is, that yield *x*’s that we can use for various inferential purposes. When we have repeated measurements of the same items, we can use those samples to our advantage. Given 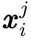, which is the j*^th^* measurement of sample *i*, we would expect 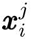 to be more similar to 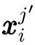 (another measurement of the same item), than to any measurement of a different item 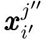. Formally, let 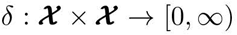 be a distance metric, we define the population `Discr`:

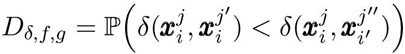

That is, “population `Discr`” *D* represents the average probability that the *within-item distance* 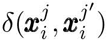 is less than the *between-item distance* 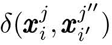. `Discr` depends on the choice of distance *δ*, as well as the measurement protocal *f* and the analysis choices *g*.

The population `Discr` represents a property of the distribution of ***θ**_i_*. In real data since we do not observe the true distribution, we instead rely on the sample `Discr`. Suppose a dataset consists of *i* ∈ {1, … , *n*} items, where each item *i* has *J_i_* repeat measurements. The sample `Discr` is defined:

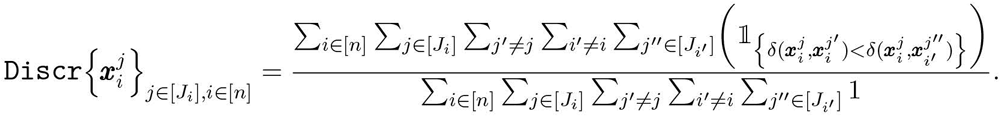

It can be shown [1] that the under the multivariate additive noise model in Assumption 1, that the sample `Discr` is both a consistent and unbiased estimator for population `Discr`.

## Supporting Information 3

### S3 Discr Provides an Informative Bound for Inference

During experimental design, the extent of subsequent inference tasks may be unknown. A natural question may be, what are the implications of the selection of a discriminable experimental design? Formally, assume the task of interest is binary classification: that is, 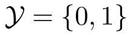, and we seek a classifier 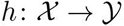. The goal of experimental design in this context is to choose the options (*f*\*, *g*\*) that will minimize the classification loss:

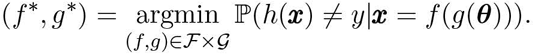

For a fixed (*f, g*), the minimal prediction error is achieved by the Bayes optimal classifier [1]:

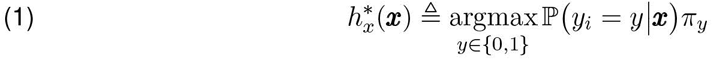

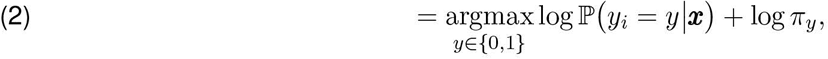

where 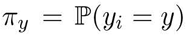, and let 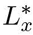 denote the error of the Bayes optimal classifier; that is, the error achieved by 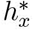.

#### Assumption 1 Multivariate Additive Noise Setting

The multivariate additive noise setting can be described as follows. For items *i* = 1, … , *n* and sessions *j* = 1, … , *s*:

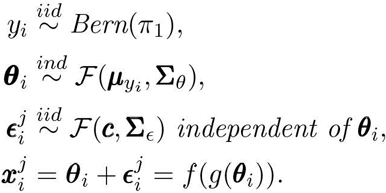

where 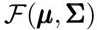 denotes a distribution with a finite mean vector **μ** and a finite, non-singular covariance **Σ**.

To connect the above model more directly with Eq. (1), we can let look at a special case

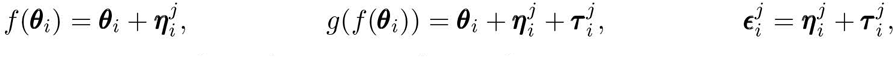

where we assume that 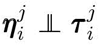, and both 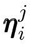 and 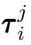 are multivariate Gaussian. Using Bayes rule and Assumption 1, note that the probability that an observation 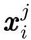 is from class *y* is given by:

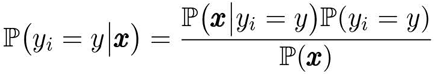

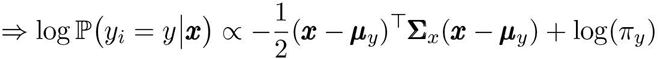

where **Σ**_*x*_ = **Σ**_*θ*_ + **Σ**_*ϵ*_ is constant between the two classes (that is, the variance is homoscedastic), and *y* is a generic value in {0, 1} that a realization *y_i_* can take. This follows directly by taking the log of the density function of the multivariate normal distribution, and removing terms not proportional in *y*. The Bayes optimal classifier is:

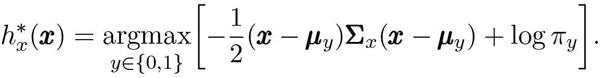

In the general case, the Bayes optimal error can be computed explicitly using that:

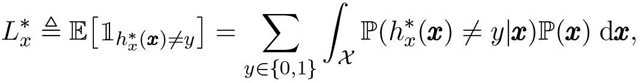

using standard rules of integration. Even when the true class distributions are known, however, computation of this integral explicitly tends to be rather tedious. For this reason, much work is dedicated to identifying cases in which the Bayes error can be bounded.

Importantly, the Bayes error can, in fact, be upper bounded by a decreasing function of `Discr`, as shown in the theorem below. In words, this theorem specifies the desirability of high `Discr`: a higher discriminability results in a lower bound on the error of future inferential tasks. Correspondingly, a strategy with a higher discriminability will have a lower bound on the error than another strategy with a lower discriminability.

#### Theorem 2.

Let 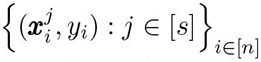 follow the multivariate additive noise setting, given in Assumption 1. Then there exists a decreasing function γ(·) of the discriminability D where:

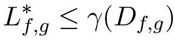

where L* is the Bayes error, or the error achieved by the Bayes optimal classifier 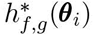.

#### Proof of Theorem (2).

Consider the additive noise setting, that is 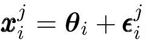,

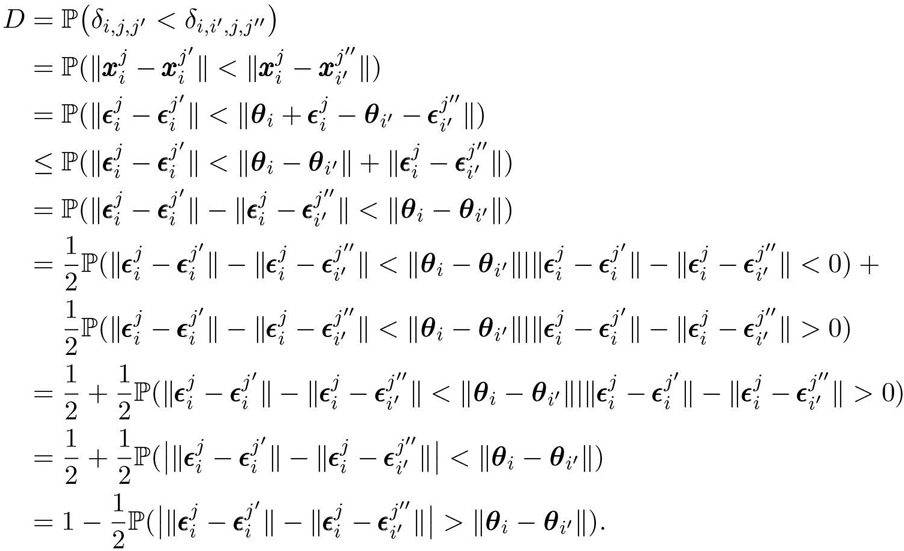

To bound the probability above, we bound the ||***θ**_i_ − **θ**_i′_*|| and 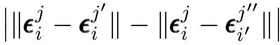 separately. We start with the first term

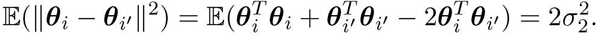

Here, 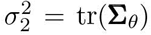 is the trace of covariance matrix of ***θ**_i_*. We can apply Markov’s Inequality for any *t* > 0:

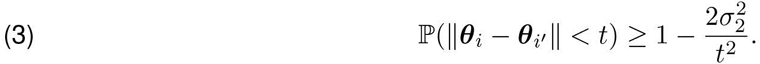

Let *a* and *b* be two constants satisfying:

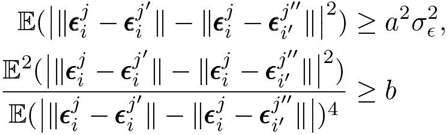

Furthermore, let 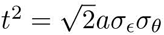 and define:

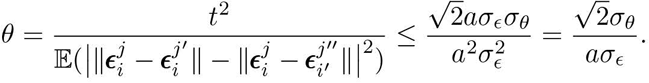

If 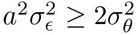, then *θ* ≤ 1. According to the Paley-Zygmund Inequality [2], that is:

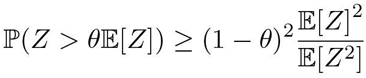

for all 0 ≤ *θ* ≤ 1 and *Z* ≥ 0, we can plug in the *θ* above to achieve

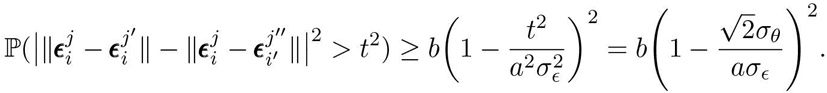

Plugging *t*^2^ into the inequality in Equation (3), we have:

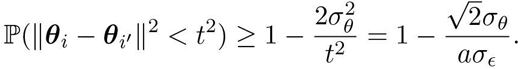

Given that ***θ**_i_*’s and 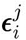 are independent by supposition, we can combine the two inequalities:

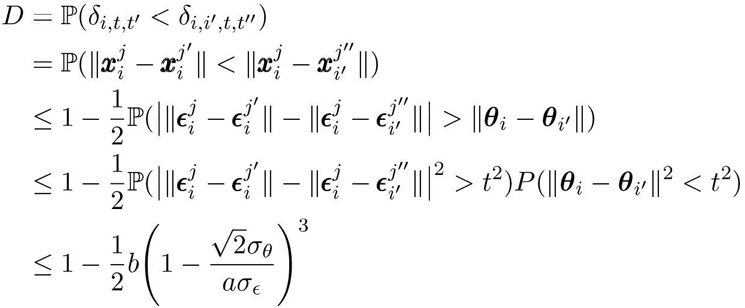

Note that the resulted bound holds true even if 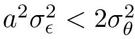, as the right hand side becomes greater than 1. This produces a bound for 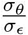:

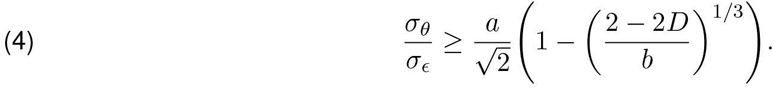

To obtain a bound on Bayes error, we use the following two observations:

1. The weighted covariance matrix of the measurements is non-singular: Define **Σ**_*x*_ as the weighted covariance matrix of ***x***:

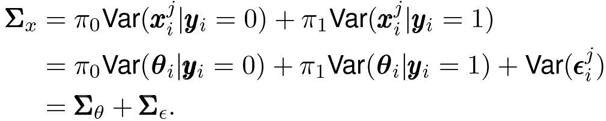

which follows since *π*_0_ +*π*_1_ = 1. Further, note that since both **Σ**_***θ***_ and **Σ**_*ϵ*_ are finite and non-singular, their sum **Σ**_*x*_ is also finite and non-singular.
2. The between-class difference is finite: Denote ∆***μ*** to be the difference between the means of the two classes. Since 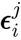 is assumed to be independent of ***y**_i_*:

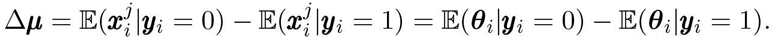

We apply Devijver and Kittler’s result [3], from equation (2.93), which gives that:

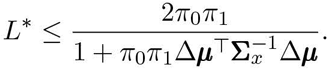

Denote 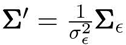. By inequality (4), note that 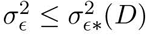, where:

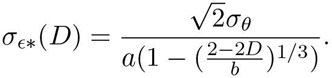

Hence, 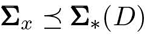 where:

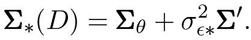

Therefore, 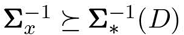, and we obtain:

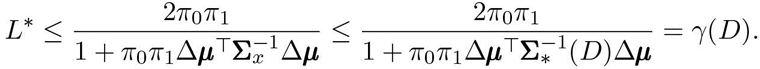

where 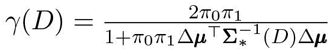 is decreasing in *D*.

Next, we will generalize this theorem to a broader class of stochastic measurements. A local ordinal embedding [4] 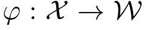 with respect to a pair of distance metrics *δ_x_, δ_w_* for a set of measurements 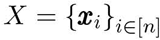 is defined as a function where if ***x**_i_, **x**_i′_, **x**_j_, **x**_j′_* ∈ *X*, then:

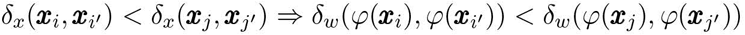

Effectively, the statement asserts that if a pair of points are closer than another pair of points, than the pair of embedded points are closer than the other pair of embedded points. In other words, the *ordering of distances* is preserved after embedding with *φ*. While this fairly broad class of embeddings preserves discriminability rather trivially, in fact, an even broader class embeddings will further preserve discriminability. In particular, an embedding need only preserve *within-item* distance orderings, rather than *all pairs* of distances. We define this class of embeddings as a **within-item** ordinal embedding. Suppose that 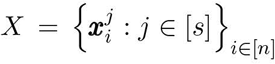 denotes a set of measurements of *n* individuals, measured *s* times each. If 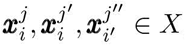, then:

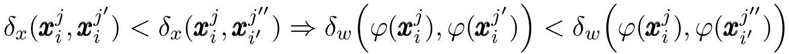

This class of embeddings instead need only preserve within-item distance relationships. Note that 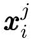 and 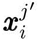 are two different measurements of the same item, and 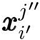 is an arbitrary measurement from a different item. If 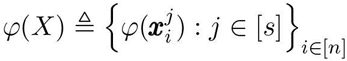 is the set of points embedded by the within-item ordinal embedding *φ*, then the discriminability of *φ*(*X*) is clearly the same as the discriminability of *X*. This is because the statement of a within-item ordinal embedding asserts that the relationship specified by discriminability holds *absolutely* (and therefore, it certainly also holds in probability). Note further that the class of embeddings which are local ordinal embeddings are a subset of the class of embeddings which are within-item ordinal embeddings.

Further, note that if *φ* were one-to-one, that the Bayes error is the same, which can be seen through a change of variables argument. These observations motivate the following corollary:

#### Corollary 3.

Suppose that 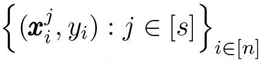 are stochastic measurements and class labels following the additive gaussian noise setting, described in Assumption 1.

Let 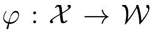 be a within-item ordinal embedding which is also one-to-one, and denote 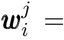 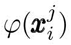. There exists a decreasing function γ(·) of the discriminability 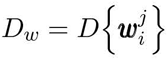 where:

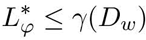

*Proof.* Denote *γ_x_*(·) to be the decreasing function of 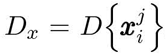 which exists by Theorem (2), where:

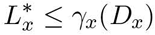

Let 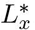 be the Bayes’ error of 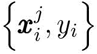. We note the following two facts:

1. The Bayes error 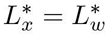: Follows since *φ* is one-to-one.
2. *D_x_* = *D_w_*: Follows since *φ* is a local ordinal embedding.

Finally, using these two facts, note that:

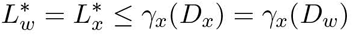

So selecting the same function *γ* = *γ_x_* gives a function of the discriminability of 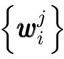 which upper bounds the Bayes’ error of 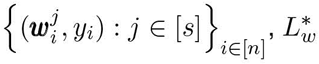, as desired.

#### Corollary 4.

Assume (f_1_, g_1_) and (f_2_, g_2_) are two analysis strategies, and suppose that 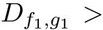 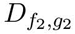. Then the bound on the Bayes error for (f_1_, g_1_) is lower than the bound on the Bayes error on (f_2_, g_2_).

*Proof.* Direct application of Theorem 2, noting that 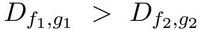 implies that 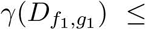 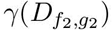 since *γ* is decreasing in *D*.

Consequently, under the described setting, the pipeline that achieves a higher `Discr` has a lower bound on the Bayes error than competing strategies, despite the fact that the task is unknown during data acquisition and analysis. Complementarily, note that if we were to instead consider the predictive accuracy 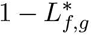, we can obtain a similar result to obtain a lower bound on the predictive accuracy via an increasing function of `Discr`. That is, in the context of the corollary, a more discriminable pipeline will tend to have a higher bound on the accuracy for an arbitrary predictive task.

## Supporting Information 4

### S4 Simulations

The following simulations were constructed, where *σ_min_*, *σ_max_* are the variance ranges, and settings were run at 15 intervals in [*σ_min_, σ_max_*] for 500 repetitions per setting. For a simulation setting with variance *σ*, the variance is reported as the normalized variance, 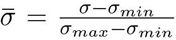. Dimensionality is 2, the number of items is *K*, and the total number of measurements across all items is 128. Typically, *i* indicates the individual identifier, and *j* the measurement index. Notationally, in the below descriptions, we adopt the convention that 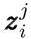 obeys the true distribution for a single observation *j* of item *i*, and 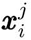 incorporates the controlled error term 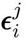, which is the term which is varied the simulation. Further, each item features 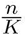 measurements.

#### Goodness of Fit Testing and Bayes Error

1. No Signal: *K* = 2 items, where the true distributions for class 1 and class 2 are the same.

- 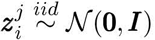, *i* = 1, … , 2, *t* = 1, … , 64. Note: **0** ∈ℝ^2^ is **0**, and likewise for ***I***
- 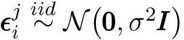, *σ* ∈ [0, 20]
- 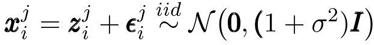
2. Cross: *K* = 2 items, where the true distributions for class 1 and class 2 are orthogonal.

- 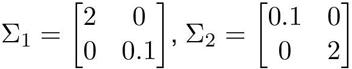
- 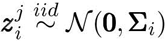 *i* = 1, 2
- 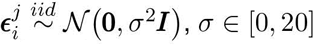
- 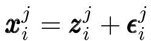
3. Gaussian: *K* = 16 items, where the true distributions are each gaussian.

- 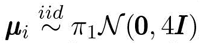 *i* = 1, … , 16
- 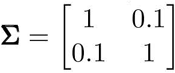
- 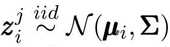
- 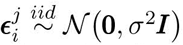, *σ* ∈ [0, 20]
- 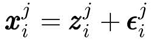
4. Ball/Circle: *K* = 2 items, where 1 item is uniformly distributed on the unit ball with gaussian error, and the second item is uniformly distributed on the unit sphere with gaussian error.

- 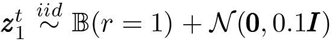 samples uniformly on unit ball of radius 2 with Gaussian error
- 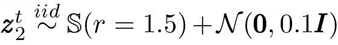 samples uniformly on unit sphere of radius 2 with Gaussian error
- 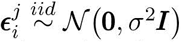, *σ* ∈ [0, 10]
- 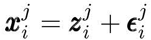
5. XOR: *K* = 2 items, where:

- 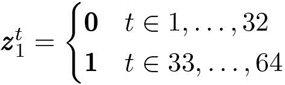
- 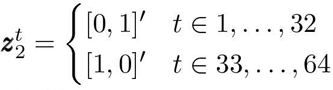
- 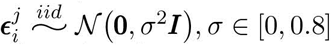
- 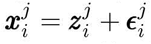

Bayes error was estimated by simulating *n* = 10,000 points according to the above simulation settings, and approximating the Bayes error through numerical integration. The classification labels for *K* = 2 simulations were consistent with the individual labels, and for the *K* = 16, the first class consists of the 8 distributions whose means were leftmost, and the rest of the distributions were the other class.

#### Comparison Testing

Items are sampled with the same true distributions 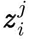 as before, with the following augmentation:

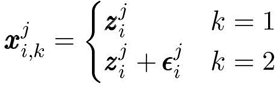

That is, the observed data 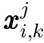 for item *i*, observation *j*, and sample *k* ∈ [2] is such that the first sample is distributed according to the true item distribution, and the second sample is distributed according to the true item distribution with an added noise term, where 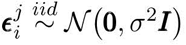:

1. No Signal: *K* = 2 *σ* ∈ [0, 10]
2. Cross: *K* = 2 *σ* ∈ [0, 1]
3. Gaussian: *K* = 16 *σ* ∈ [0, 1]
4. Ball/Circle: *K* = 2 *σ* ∈ [0, 1]
5. XOR: *K* = 2 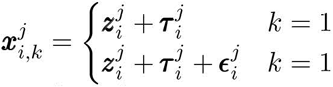 where 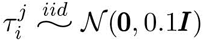 *σ* ∈ [0, 0.2]

By construction, one would anticipate `Discr` of the first sample to exceed that of the second sample, as the second sample has additional error. Therefore, the natural hypothesis is:

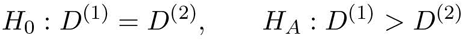

## Supporting Information 5

### S5 Hypothesis Testing

#### Goodness of Fit Test

Recall the goodness of fit test, shown in Equation (1). We approximate the distribution of 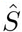 under the null through a permutation approach. The item labels of our *N* samples are first permutated randomly, and 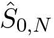 is computed each time given the observed data 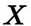 and the permuted labels. For a level *α* significance test, we compare 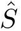 to the (1 − *α*) quantile 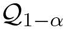 of the empirical null distribution 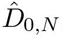, and reject the null hypothesis if 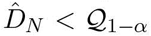. This approach provides a consistent and valid test under general assumptions.

Note that the permutation-based approach requires *r* computations of the sample `Discr`. The total computational complexity is then 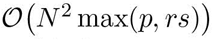. This approach is only linear in the number of desired repetitions, and therefore is sensible for most settings in which the sample `Discr` can itself be computed. Moreover, we can greatly speed this computation up through parallelization. With *T* cores, the computational complexity is instead 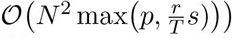, as shown in Algorithm 1. We extend this goodness of fit test to both `PICC` and `I2C2` to provide a robust *p*-value associated with both statistics of interest. Note that the permutation approach can be generalized to any statistic quantifying replicability based on repeated measurements.

**Algorithm 1.**
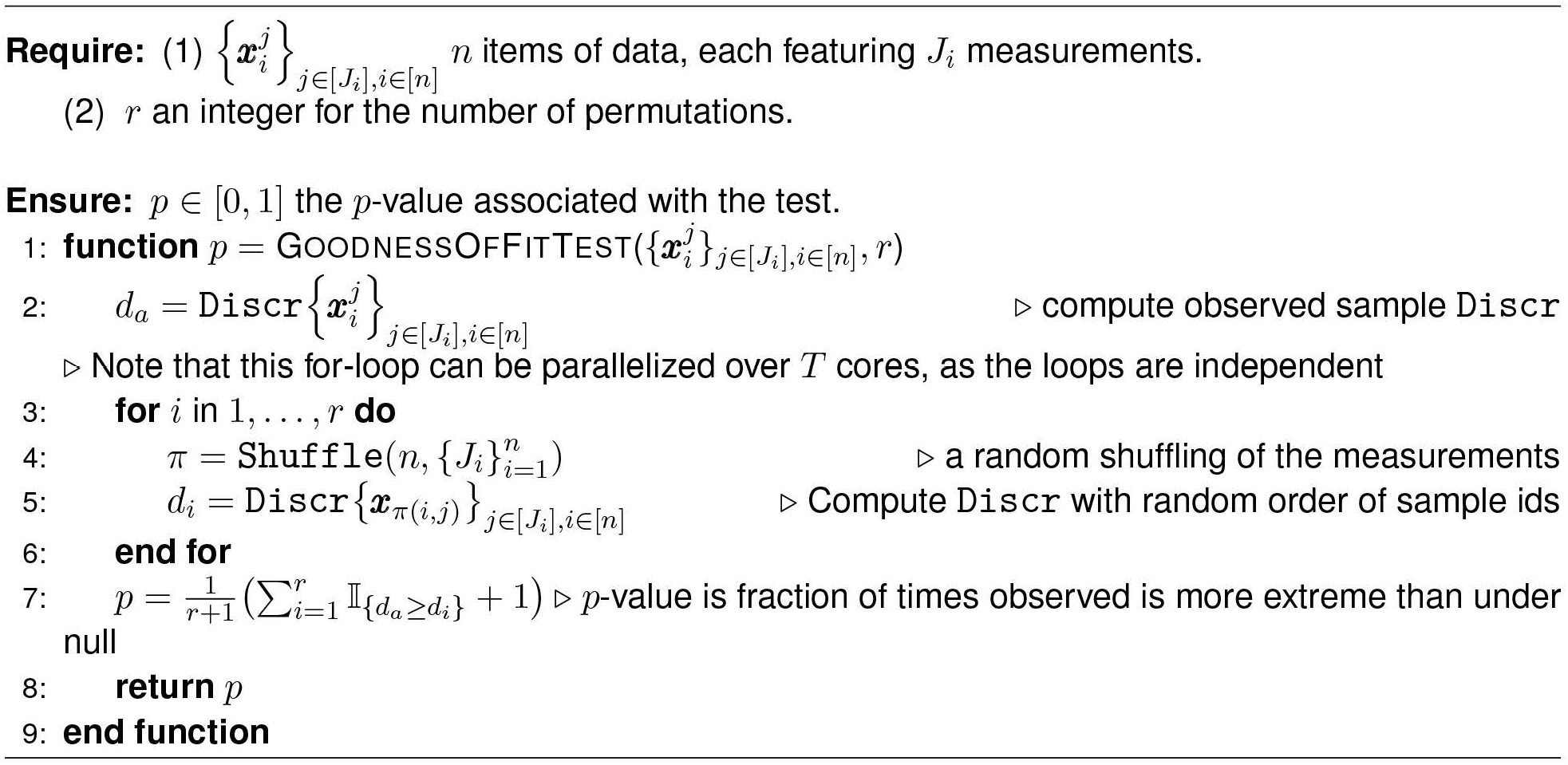
Discr Goodness of Fit Test. Our implementation of the permutation test for the goodness of fit test of the hypothesis given in Equation (1) requires 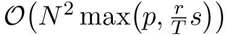 time, where *r* is the number of permutations and *T* is the number of cores available for the permutation test. The `Shuffle` function is the function which rearranges all of the data within the dataset, without regard to item nor measurement index. The output provides a new measurement index for each item *i* and measurement *j*.

#### Comparison Test

We implement Comparison testing using a permutation approach, similar to the goodness of fit test. First, compute the observed difference in `Discr` between two design choices. The null distribution of the difference in `Discr` is constructed by first taking random convex combinations of the observed data from each of the two methods choices (the “randomly combined datasets”). `Discr` is computed for each of the two randomly combined datasets for each permutation. Finally, for each permutation, the all pairs of observed differences in `Discr` is computed. Finally, the observed statistic is compared with the differences under the null of the randomly combined datasets. The p-value is the fraction of times that the observed statistic is more extreme than the null. Note that we can use this approach for both one and two-tailed hypotheses for an experimental design having higher `Discr`, lower `Discr`, and equal `Discr` relative a second approach; we implement all three in the software implementation of the comparison test. The Algorithm for the comparison test is shown in Algorithm 2, with the alternative hypothesis as specified in Equation (2). The computational complexity is then 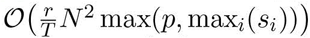. Note that for each permutation, the limiting step is the computation of the `Discr` in 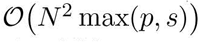. This is then offset through parallelization over *T* cores in the implementation. We extend this comparison test to all competing approaches to provide a robust *p*-value associated with both statistics of interest, for similar reasons to the above. Again, this permutation approach can be generalized to any statistic quantifying replicability based on repeated measurements.

**Algorithm 2.**
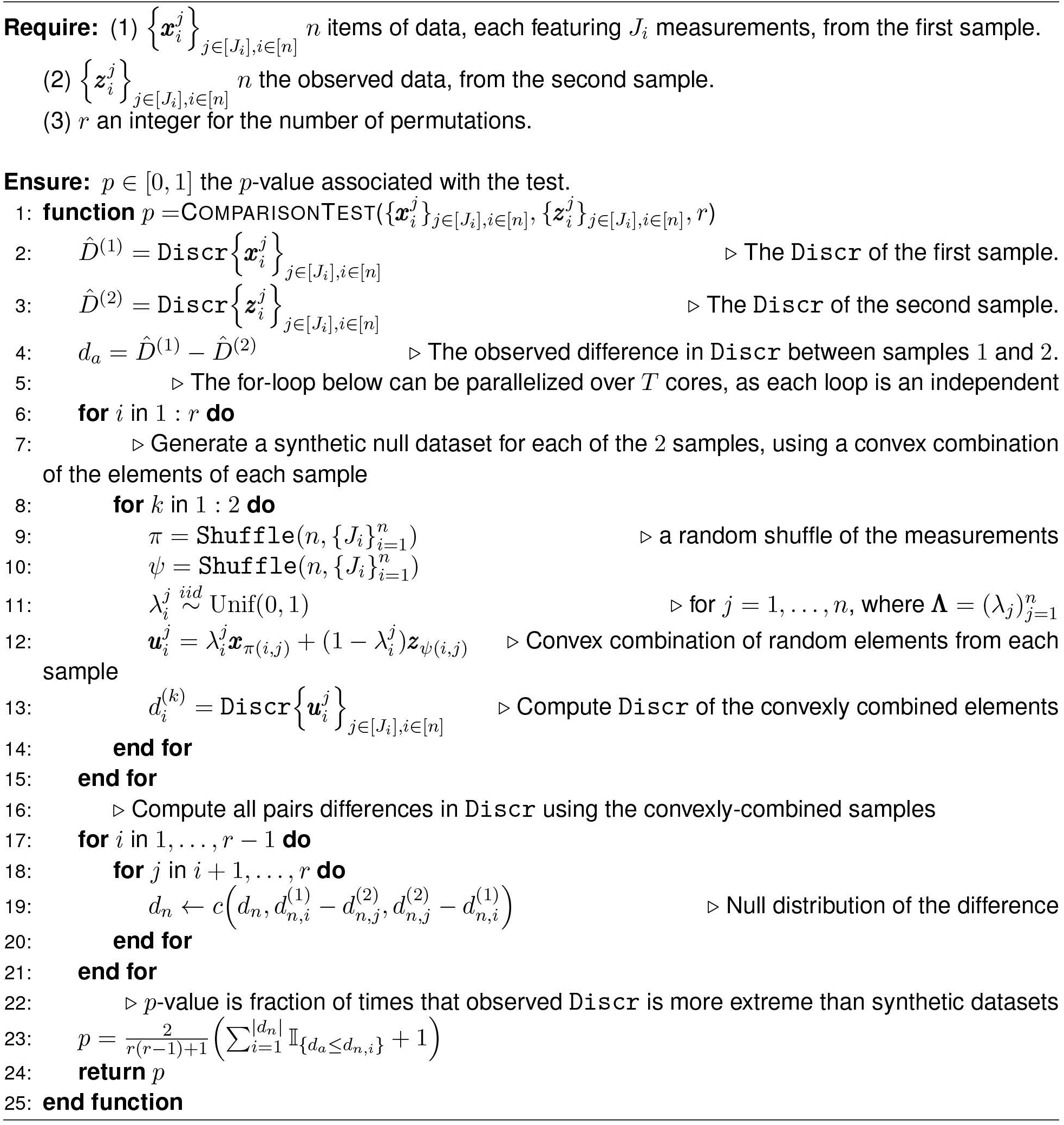
Discr Discriminability Comparison Test. Our implementation of the permutation test for the hypothesis given in Equation (2) requires 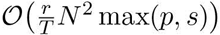 time, where *r* is the number of permutations and *T* is the number of cores available for the permutation test. Above, the only alternative considered is that *H_A_*: *D*^(1)^ > *D*^(2)^; our code-based implementation provides strategies for *H_A_*: *D*^(1)^ < *D*^(2)^ and *HA*: *D*^(1)^ = *D*^(2)^ as well.

## Supporting Information 6

### S6 Connectomics Application

#### Data Acquisition and Analysis

##### fMRI Analysis Pipelines

The fMRI connectomes were acquired as follows. Motion correction is performed via `mcflirt` to estimate the 6 motion parameters (*x*, *y*, *z* translation and rotations). Registration is performed by first performing a cross-modality registration from the functional to the anatomical MRI using `flirt-bbr`, followed by registration to the anatomical template using either (1) FSL-`fnirt` or (2) ANTs-`SyN`, two techniques for non-linear registration. Frequency filtering was performed by either not frequency filtering, or (2) bandpass filtering signal outside of the [.01, .1] Hz range. Volumes were either (1) not scrubbed, or (2) scrubbed if motion exceeded 0.5 mm, in which case the preceding volume and succeeding two volumes were removed. Global signal regression was either (1) not performed, or performed by removing the global mean signal across all voxels in the functional timeseries. Moreover, across all analysis pipelines, the top 5 principal components (`compcor`), Friston 24 parameters, and a quadratic polynomial were fit and regressed from the functional timeseries. Finally, the voxelwise timeseries were spatially downsampled using (1) the CC200 parcellation, (2) the AAL parcellation, (3) the Harvard-Oxford parcellation, or (4) the Desikan-Killany parcellation. Graphs were estimated by (1) computing the rank of the non-zero raw absolute correlations (zero-weight edges given a value of 0), log-transforming the raw absolute correlations (the minimum value of the graph is down-scaled by a factor of 100 and then added to each edge to eliminate taking log of zero-weight edges), or (3) computing the raw absolute correlation between pairs of regions of interest in each parcellation. No mean centering was performed for functional connectivity estimates. Specific data analysis instructions for deployment in `AWS` can be found in the https://neurodata.io/m2g. All data analysis was performed in the `AWS` cloud using `CPAC` version `3.9.2` [1]. All parcellations are available in `neuroparc` human brain atlases [2].

##### dMRI Analysis Pipelines

The dMRI connectomes were acquired as follows. The dMRI scans were corrected for eddy currents using FSL’s `eddy-correct` [3]. FSL’s “standard” linear registration pipeline was used to register the sMRI and dMRI images to the MNI152 atlas [3–6]. A tensor model is fit using DiPy [7] to obtain an estimated tensor at each voxel. A deterministic tractography algorithm is applied using DiPy’s EuDX [7, 8] to obtain streamlines, which indicate the voxels connected by an axonal fiber tract. Graphs are formed by contracting voxels into graph vertices depending on spatial [9], anatomical [10–13], or functional [14–17] similarity. Given a parcellation with vertices *V* and a corresponding mapping *P*(*u*) indicating the voxels within a region *i*, we contract our fiber streamlines as follows. 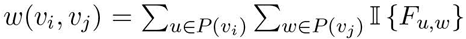 where *F_u,w_* is true if a fiber tract exists between voxels *u* and *w*, and false if there is no fiber tract between voxels *u* and *w*. The specific parcellations leveraged are detailed in **(author?)** [18], consisting of parcellations defined in the MNI152 space [10–17]. The graphs are then re-weighted using the afforementioned weighting schemes described in fMRI Analysis Pipelines Supplementary Information; namely, the raw, ranked, and log edge-weights. All parcellations are available in `neuroparc` human brain atlases [2].

##### PCR RealSeqS Cancer Genomics Pipeline

The RealSeqS samples were acquired as follows. PCR was performed in 25 *μL* reactions containing 7.25 *μL* of water, 0.125 *μL* of each primer, 12.5 *μL* of NEBNext Ultra II Q5 Master Mix (New England Biolabs cat # M0544S), and 5 *μL* of DNA. The cycling conditions were: one cycle of 98*°*C for 120 s, then 15 cycles of 98*°*C for 10 s, 57*°*C for 120 s, and 72*°*C for 120 s. Each plasma DNA sample was assessed in eight independent reactions, and the amount of DNA per reaction varied from 0.1 *μg* to 0.25 *μg*. A second round of PCR was then performed to add dual indexes (barcodes) to each PCR product prior to sequencing. The second round of PCR was performed in 25 *μL* reactions containing 7.25 *μL* of water, 0.125 *μL* of each primer, 12.5 *μL* of NEBNext Ultra II Q5 Master Mix (New England Biolabs cat # M0544S), and 5 uL of DNA containing 5% of the PCR product from the first round. The cycling conditions were: one cycle of 9 8ÂřrC for 120 s, then 15 cycles of 98*°*C for 10 s, 65*°*C for 15 s, and 72*°*C for 120 s. Amplification products from the second round were purified with AMPure XP beads (Beckman cat # a 63880), as per the manufacturer’s instructions, prior to sequencing. As noted above, each sample was amplified in eight independent PCRs in the first round. Each of the eight independent PCRs was then re-amplified using index primers in the second PCR round. Bowite2 was then used to align reads to the human reference genome assembly GRC37 [19] for each well. After alignment to ∼ 750,000 amplicons, the wells were downsampled into non-overlapping windows of 5 × 10^4^ bases, 5 × 10^5^ bases, 5 × 10^6^ bases, or to the individual chromosome level (the resolution of the data).

#### Effect Size Investigation

In this investigation, we are interested in learning how maximization based on the observed notion of replicability correlates with real performance on a downstream inference task. Recalling Corollary 4 from S3, we explore the implications of this corollary in a large neuroimaging dataset provided by the Consortium for Reliability and Reproducibility [20], and demonstrate that selection of the experimental design via `Discr`, in fact, facilitates improved downstream inference on both a regression and classification task. We further extend this to two separate genomics datasets investigating classification tasks, and again demonstrate that selection of experimental design via `Discr` improves downstream inference. This provides strong motivation for leveraging the `Discr` for experimental design.

Ideally, for a particular summary reference statistic, a high value will generally correlate with a positive effect size. For datasets *i* = 1, … , *M* where *M* is the total number of datasets, an analysis strategy *j* = 1, … , 192 for 192 total analysis strategies, and *k* = 1, … , 3 are our summary reference statistics of interest (`Discr, PICC, Fingerprint, I2C2, Kernel`), we fit the standard linear regression model *Y* = *βX* + *∊*, where we model the effect size *Y* estimated by `DCorr` [21] via a linear relationship with *X*, the observed reference statistic for approach *k*, with coefficient *β*. Note that the interpretation of *β* is the expected change in the effect size *Y* due to a single unit change in the observed reference statistic *X*. Both *Y* and *X* are uniformly normalized across all strategies within a single dataset to facilitate intuitive comparison across methods. For each reference statistic *k*, we pose the following hypothesis:

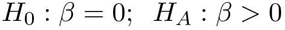

Acceptance of the alternative hypothesis would have the interpretation that an increase in the observed reference statistic *X* would tend to correspond to an increase in the observed effect size *Y* , and the relevant test is the one-way *Z*-test. To robustify against model assumptions, we use robust standard errors [22]. Acceptance of the alternative hypothesis against the null provides evidence that an increase in the sample statistic corresponds to an increase in the observed effect size, where the responses (age, sex, cancer status) were not considered at the time the data were analyzed nor when the reference statistics computed. This provides evidence that the statistic is informative for experimental design within the context of this investigation. Model fitting for this investigation is conducted using the `lm` package in the `R` programming language [23].

**S6 Table 1.** fMRI Dataset Descriptions. In the above table, `STC` corresponds to slice timing correction. Rows with NA entries do not have available metadata associated with the scanning protocol. The column TRT indicates the follow up time for retest. A value of None indicates that the scans were back to back. The sample `Discr` corresponds to the `Discr` of the best performing pipeline overall, `FNNNCP`. *The test-retest structure for NYU1 was 5 - 11 months between sessions 1 and 2, and 30 − 45 minutes between sessions 2 and 3.

| Dataset | Manuf. | Model | TE (ms) | TR (ms) | STC | #Timepts | #Sub | #Ses | #Scans | TRT (days) | Discr |
| --- | --- | --- | --- | --- | --- | --- | --- | --- | --- | --- | --- |
| KKI2009 | Philips | Achieva | 30 | 2000 | seq. | 210 | 21 | 2 | 42 | <1 | 0.93 |
| NKI24 | Siemens | TrioTim | 30 | 645 | inter. | 900 | 24 | 2 | 47 | <14 | 0.98 |
| BNU1 | Siemens | TrioTim | 30 | 2000 | inter. | 200 | 50 | 2 | 100 | 42 | 0.97 |
| BNU2 | Siemens | TrioTim | 30 | variable | inter. | variable | 50 | 2 | 100 | 103 | 0.92 |
| DC1 | Philips | NA | 35 | 2500 | inter. | 120 | 114 | 4 | 244 | ? | 0.95 |
| HNU1 | GE | MR750 | 30 | 2000 | inter. | 300 | 30 | 10 | 300 | 3 | 0.98 |
| IACAS | GE | Signa | 30 | 2000 | inter. | 240 | 28 | 3 | 59 | 42 | 0.83 |
| IBATRT | Siemens | TrioTim | 30 | 1750 | seq. | 220 | 36 | 2 | 50 | None | 0.95 |
| IPCAS | NA | NA | NA | NA | NA | NA | 78 | 2 | 156 | – | 0.99 |
| IPCAS1 | Siemens | TrioTim | 30 | 2000 | inter. | 205 | 30 | 2 | 60 | 7 | 1.00 |
| IPCAS2 | Siemens | TrioTim | 30 | 2500 | inter. | 212 | 35 | 2 | 70 | 30 | 0.98 |
| IPCAS5 | Siemens | TrioTim | 30 | 2000 | inter. | 170 | 22 | 2 | 44 | >10 (min) | 0.96 |
| IPCAS6 | Siemens | TrioTim | 30 | 2500 | inter. | 242 | 2 | 15 | 30 | 3 (hrs) | 1.00 |
| IPCAS8 | Siemens | TrioTim | 30 | 2000 | inter. | 240 | 13 | 2 | 26 | >1 (years) | 0.96 |
| JHNU | Siemens | TrioTim | 30 | 2000 | inter. | 250 | 30 | 2 | 60 | NA | 0.96 |
| LMU3 | Siemens | TrioTim | 30 | 3000 | inter. | 120 | 25 | 2 | 50 | NA | 0.93 |
| MRN1 | NA | NA | NA | NA | NA | NA | 53 | 2 | 88 | 120 | 0.94 |
| NYU1 | Siemens | Allegra | 25 | 2000 | NaN | 197 | 25 | 3 | 75 | 5-11 (mo)* | 0.98 |
| NYU2 | Siemens | Allegra | 15 | 2000 | inter. | 180 | 187 | 3 | 252 | <1 (hrs) | 0.96 |
| SWU1 | Siemens | TrioTim | 30 | 2000 | inter. | 240 | 20 | 3 | 59 | NA | 0.97 |
| SWU2 | Siemens | TrioTim | 30 | 2000 | inter. | 300 | 27 | 2 | 54 | NA | 0.96 |
| SWU3 | Siemens | TrioTim | 30 | 2000 | inter. | 242 | 24 | 2 | 48 | NA | 0.98 |
| SWU4 | Siemens | TrioTim | 30 | 2000 | inter. | 242 | 235 | 2 | 467 | 1 (yrs) | 0.97 |
| UM | Siemens | TrioTim | 30 | 2000 | seq. | 150 | 80 | 2 | 160 | NA | 0.99 |
| UPSM1 | Siemens | TrioTim | 29 | 1500 | seq. | 200 | 100 | 3 | 230 | 473 - 1434 | 0.89 |
| Utah1 | Siemens | TrioTim | 28 | 2000 | inter. | 240 | 26 | 2 | 52 | >2 (yrs) | 0.92 |
| UWM | GE | MR750 | 25 | 2600 | inter. | 231 | 25 | 2 | 50 | NA | 0.96 |
| XHCUMS | Siemens | TrioTim | 30 | 3000 | inter. | 124 | 24 | 5 | 120 | 180 | 0.91 |

#### Human Brain Imaging Dataset Descriptions

**S6 Table 2.** dMRI Dataset Descriptions. In the above table, #`Dir` corresponds to the number of diffusion directions. Rows with `NA` entries do not have available metadata associated with the scanning protocol. The sample `Discr` corresponds to the `Discr` of the pipeline with the `CPAC200` parcellation and the log-transformed edges.

| Dataset | Manuf. | Model | TE (ms) | TR (ms) | #Dir | bval $\frac{s}{mm^2}$ | #Sub | #Ses | #Scans | TRT (days) | Discr |
| --- | --- | --- | --- | --- | --- | --- | --- | --- | --- | --- | --- |
| BNU1 | Siemens | TrioTim | 89 | 8000 | 30 | 1000 | 57 | 2 | 113 | 42 | 1.00 |
| HNU1 | GE | MR750 | Min | 8600 | 33 | 1000 | 30 | 10 | 300 | 3 | 0.99 |
| KKI2009 | Philips | NA | 32 | 6281 | 65 | 700 | 21 | 2 | 42 | <1 | 1.00 |
| NKI24 | Siemens | TrioTim | 95 | 2400 | 137 | 1500 | 20 | 2 | 40 | <14 | 1.00 |
| SWU4 | Siemens | TrioTim | NaN | NaN | 93 | 1000 | 227 | 2 | 454 | 1 (yrs) | 0.88 |

##### Useful Data Links

All relevant analysis scripts and data for figure reproduction in this manuscript made publicly available, and can be found at https://neurodata.io/mgc.

## Supporting Information 7

### S7 Discriminability Decomposition

Consider data which is observed as the pairs 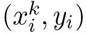, where *i* = 1, … , *n* indexes subjects, and *k* = 1, … , *s* indexes sessions. We suppose that 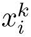 represents a measurement of interest, and *y_i_* represents a subject-specific categorical class of interest (such as a natively categorical covariate such as sex, or a natively numeric covariate such as age which can be coerced to categorical; e.g., using age quintiles or deciles). Interestingly, the discriminability can be separated into the within-class and between-class contributions on the basis of *y_i_*.

#### Within-Class Discriminability

Let 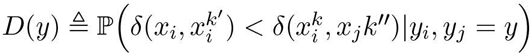 be the discriminability for class *y* Note that:

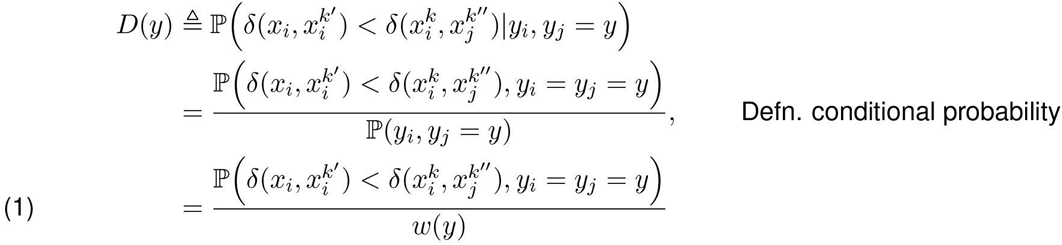

where we define 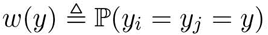.

Consider the within-class discriminability 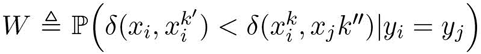. This quantity can be interpreted as the discriminability, conditional on two items being from the same class. Note that:

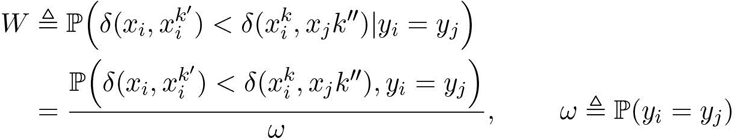

By the law of total probability, note that:

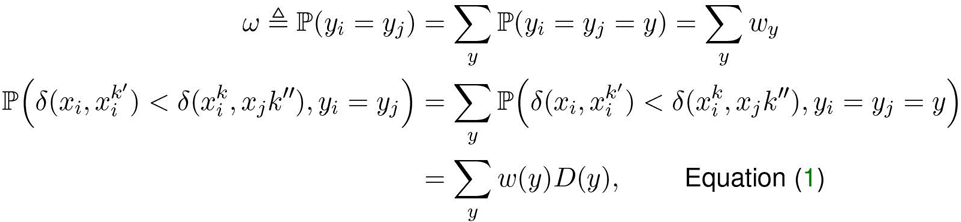

Which shows that:

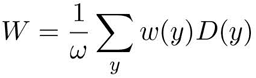

or that the within-class discriminability is a weighted sum of the per-class discriminabilities *D*(*y*), weighted by the probability of a pair of items being in class *y*.

#### Between-Class Discriminability

Let 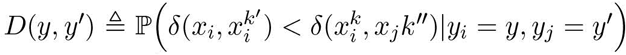 be the discriminability of items in class *y* to items in class *y*′. Note that:

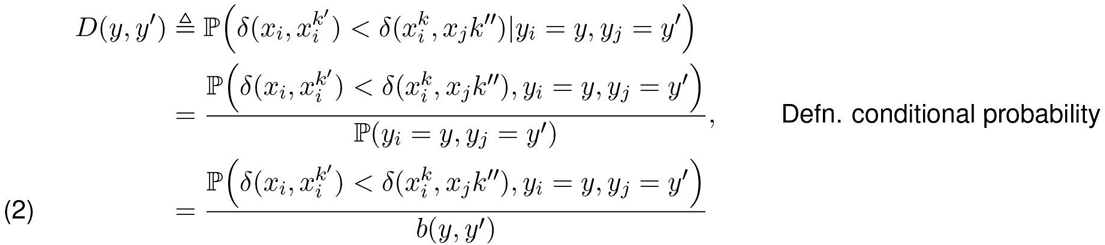

Where we define 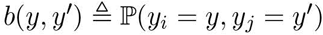.

Consider the between-class discriminability 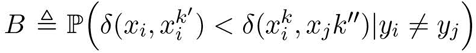. This quantity can be interpreted as the discriminability, conditional on two items being from a different class. Note that:

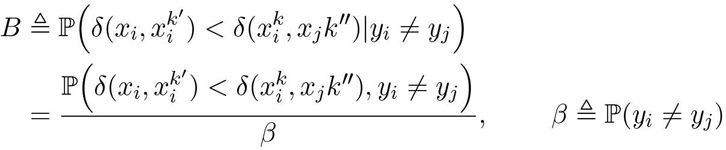

Again, using the law of total probability:

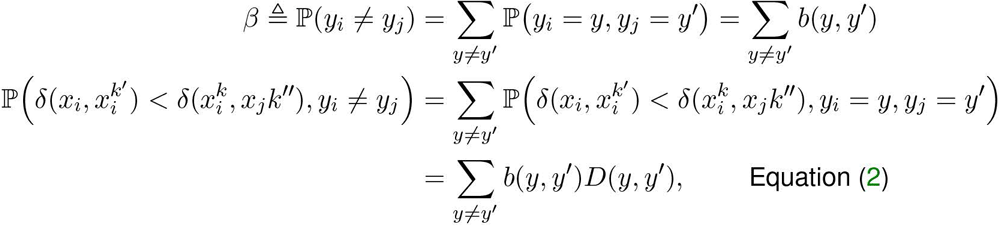

Which shows that:

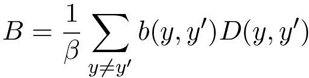

#### Discriminability Decomposition

Finally, note that:

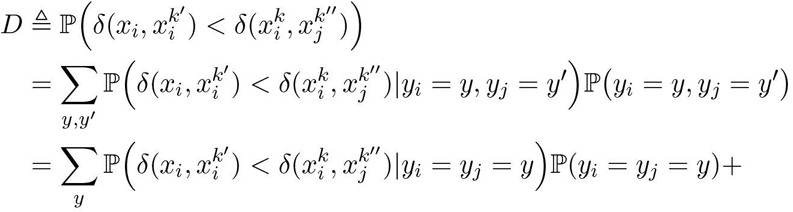

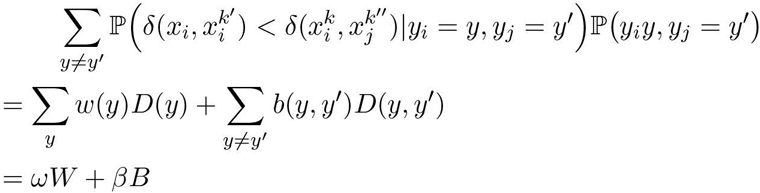

Showing that discriminability can be decomposed as a weighted sum of the within and between-class discriminabilities.

